# Molecular basis of polyadenylated RNA fate determination in the nucleus

**DOI:** 10.1101/2025.09.16.676470

**Authors:** Andrii Bugai, Ulrich Hohmann, Ana Lorenzo, Max Graf, Laura Fin, Jérôme O. Rouvière, Laszlo Tirian, Yuhui Dou, Patrik Polák, Dennis Johnsen, Lis Jakobsen, Jens Skorstengaard Andersen, Julius Brennecke, Clemens Plaschka, Torben Heick Jensen

## Abstract

Eukaryotic genomes generate a plethora of polyadenylated (pA^+^) RNAs^1,2^, that are packaged into ribonucleoprotein particles (RNPs). To ensure faithful gene expression, functional pA^+^ RNPs, including protein-coding RNPs, are exported to the cytoplasm, while transcripts within non-functional pA^+^ RNPs are degraded in the nucleus^1–4^. How cells distinguish these opposing fates remains unknown. The DExD-box ATPase UAP56/DDX39B is a central component of functional pA^+^ RNPs, promoting their docking to the nuclear pore complex (NPC)-anchored ‘transcription and export complex 2 (TREX-2)’ (ref.^5,6^), which triggers transcript release from UAP56 to facilitate export (ref.^7,8^). Here, we uncover that the ‘Poly(A) tail exosome targeting (PAXT)’ connection^9^ harbors its own TREX-2-like module, which releases pA^+^ RNAs from UAP56 for decay by the nuclear exosome. The core of this module consists of a LENG8-PCID2-SEM1 (LENG8-PS) trimer, which we show is structurally and functionally equivalent to the central GANP-PCID2-SEM1 (GANP-PS) trimer of TREX-2. Mutagenesis and transcriptomic data demonstrate that the nuclear fate of pA^+^ RNPs is governed by the contending actions of nucleoplasmic PAXT and NPC-associated TREX-2, which interpret RNA-bound UAP56 as a signal for RNA decay or export, respectively. As RNA targets of PAXT are generally short and intron-poor, we propose an overall model for pA^+^ RNP fate determination, whereby the distinct sub-nuclear localizations of PAXT and TREX-2 govern the degradation of short non-functional pA^+^ RNAs while allowing export of their longer and functional counterparts.

## INTRODUCTION

RNA polymerase II (RNAPII) extensively transcribes mammalian genomes, yielding a wide range of unadenylated and polyadenylated RNAs^1–4^. Moreover, individual transcription units (TUs), that generate standard full-length transcripts also give rise to an array of shorter isoforms^10,11^. Thus, functional RNAs are produced alongside a wealth of futile RNAPII products. While mature functional pA^+^ RNAs, such as protein-coding mRNAs, are exported from the nucleus to the cytoplasm, their non-functional counterparts are typically retained and degraded^3,4^. This is primarily achieved by the nucleoplasmic PAXT connection, consisting of a heterodimeric core of the RNA helicase MTR4 and the Zn-finger protein ZFC3H1 (ref.^9,12^). Additional, and less well-described, interactions with the nuclear pA^+^ RNA binding protein (PABPN1) and accessory factors direct transcript turnover by the 3’-5’ exonucleolytic exosome complex^13–15^. However, despite this known inventory, the biochemical basis by which PAXT distinguishes non-functional pA^+^ RNAs remains a major unresolved question.

Prior to their nuclear export, pA^+^ RNAs are packaged with proteins into pA^+^ RNPs. Central to this process is the export factor and DExD-box ATPase UAP56, also known as DDX39B, which is recruited to pA^+^ RNPs in preparation for their nuclear export^5,16^. At the nuclear envelope, the activity of the NPC-associated GANP–PS trimer of TREX-2 (ref.^6^) facilitates the release of RNA from UAP56, enabling export^7,8^. Again, how pA^+^ RNP sorting is orchestrated to favor the selected export of functional pA^+^ RNAs is unknown.

Here, we interrogate two TREX-2-like human complexes, SAC3D1-PCID2-SEM1 (SAC3D1-PS) and LENG8-PS, in which the conserved SAC3D1 and LENG8 proteins, respectively, replace GANP. The GANP-PS, SAC3D1-PS and LENG8-PS complexes are structurally similar and share the ability to release UAP56 from RNA. Notably, we show that LENG8-PS constitutes a module of PAXT, that acts on UAP56 to promote transcript turnover in contrast to the RNA export activity of TREX-2. Our findings reveal that nuclear pA^+^ RNA export and decay employ a shared biochemical mechanism to act on pA^+^ RNPs but with fundamentally different outcomes. Based on the RNA target-specificity of PAXT and its separate nuclear localization from TREX-2, we propose a general model for pA^+^ RNP fate determination.

## RESULTS

### TREX-2-like complexes release RNA from UAP56

To mediate NPC docking of export-competent pA^+^ RNPs, UAP56 binds the five subunit TREX-2 complex (GANP, PCID2, SEM1, CETN2/3, ENY2) through the TREX-2 minimal recombinant complex core (TREX-2^M^), including PCID2, SEM1, and the SAC3 domain of the scaffolding subunit GANP^7^ (Fig. 1a, left). The ability of TREX-2^M^ to release UAP56 from the pA^+^ RNP critically depends on the conserved SAC3 domain-containing ‘wedge loop’^7,8^ (Fig. 1b). Notably, similar SAC3 domains are present in the UAP56-interacting^7^ LENG8 and SAC3D1 proteins, which are broadly conserved amongst eukaryotes, but otherwise share no sequence features with GANP or each other (Fig. 1a,b). Moreover, structural modelling by AlphaFold2 (ref.^17,18^) of the SAC3 domains of LENG8 or SAC3D1, in complex with PCID2 and SEM1, displayed marked similarities to the CryoEM structure of TREX-2^M^ (ref.^7^) (Supplementary Fig. 1a). Finally, and central to the present study, LENG8 was found in immunoprecipitates of PAXT core components ZFC3H1 and MTR4 (ref.^9,19^, also see Fig. 2 below) and recently shown to interact with PCID2 and SEM1 in both human and yeast cells^20,21^. Collectively, this prompted us to investigate these TREX-2^M^-like complexes in more detail.

**Figure 1.**
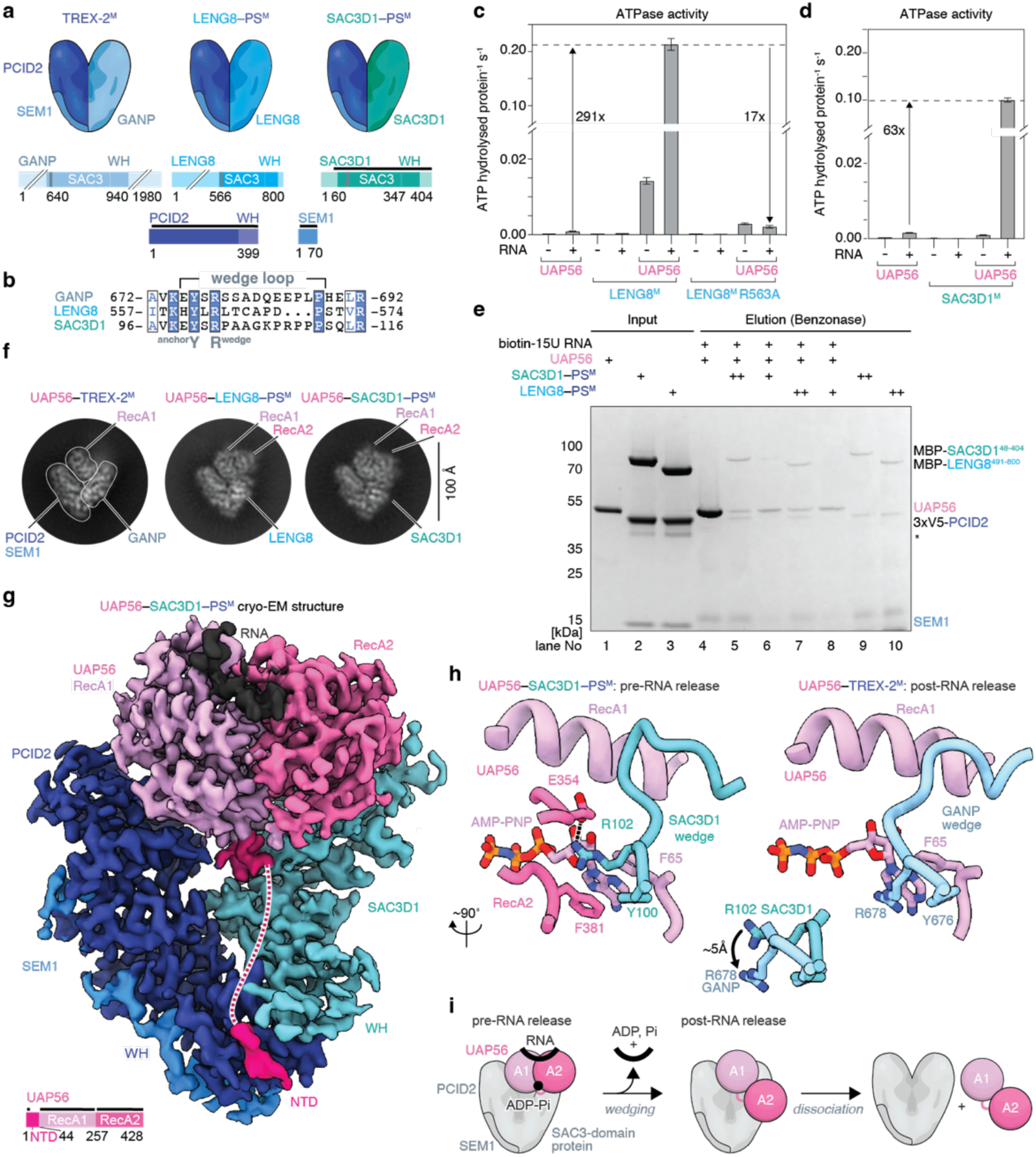
TREX-2-like complexes bind UAP56 to trigger RNA release. **a.** Cartoon models of TREX-2 and TREX-2-like core complexes with their domain architectures below: PCID2 (dark blue), SEM1 (mid blue), SAC3-domain containing proteins GANP, LENG8 and SAC3D1 (different shades of blue). Positions of conserved wedge loops inside SAC3-domains are shown as grey bars. Regions included in the atomic model in **g** are indicated by black lines. WH, Winged Helix. **b.** Multiple sequence alignment of wedge loop sequences of human GANP (Uniprot ID O60318), LENG8 (Q96PV6) and SAC3D1 (A6NKF1). Highlighted are a conserved tyrosine residue, which anchors the wedge loop on the SAC3 domain (^anchor^Y) and the central wedge loop arginine (R^wedge^). Amino acids are colored by conservation (blue text, conserved residue; blue background, invariant residue). **c-d.** ATPase assays demonstrating that LENG8–PS^M^ (**c**) and SAC3D1–PS^M^ (**d**) stimulate the apparent ATPase activity of RNA-bound UAP56. The broken Y-axes show ATPase rates (mean ± SD), the fold stimulations of which are indicated. Note presence (+) or absence (-) of a 15 poly-uridine RNA substrate. In (**c**) the reduced stimulation of UAP56’s apparent ATPase activity by the LENG8^R563A^– PS^M^ complex is shown. **e.** UAP56 release assays demonstrating the stimulatory effects of LENG8–PS^M^ and SAC3D1–PS^M^ complexes. Bead-immobilized 15 poly-uridine RNA was incubated with UAP56 and ATP to form UAP56-ADP-P_i_-RNA complexes^7^. After removal of unbound UAP56 and excess ATP, increasing amounts of recombinant LENG8-PS^M^ or SAC3D1-PS^M^ complexes were added. UAP56-ADP-P_i_-RNA complexes remaining on beads after the indicated treatments were then eluted by Benzonase digestion and analyzed by SDS-PAGE and Coomassie-staining. Asterisk denotes contaminant. **f.** Representative cryo-EM 2D classes for UAP56-TREX-2^M^ (left, ref.^7^), UAP56-LENG8–PS^M^ (middle) and UAP56-SAC3D1–PS^M^ (right), showing UAP56 in the RNA-clamped conformation, when in complex with either LENG8–PS^M^ or SAC3D1–PS^M^ but not with TREX-2^M^. **g.** UAP56-RNA-SAC3D1–PS^M^ cryo-EM density at a resolution ranging from 2.6 to 4.5 Å and colored by subunit identity. SEM1, blue; PCID2, dark blue; SAC3D1^48–404^, green blue; UAP56, shades of pink; flexible region of UAP56 N-terminal shown as dashed line; RNA, black. Cartoon at the bottom left showing UAP56 domain structure. **h.** Stick representation comparisons of details of the SAC3D1 wedge loop-UAP56-nucleotide interactions for UAP56-RNA-SAC3D1–PS^M^ (left) or UAP56-RNA-TREX-2^M^ (right). Superpositions of the wedge loop anchoring tyrosine and the central arginine of SAC3D1 or GANP are shown in the middle. GANP in light blue, colors otherwise as in panel **i.** Cartoon representation of UAP56-RNA-TREX-2/TREX-2-like complex interactions, including RNA release from UAP56. Left to right: (1) pre-release state: TREX-2/TREX-2-like complexes bind RNA-clamped UAP56; (2) post-release state: RNA is unclamped from UAP56, leaving UAP56 in an open conformation bound to the TREX-2/TREX-2-like complex; (3) dissociation.

**Figure 2.**
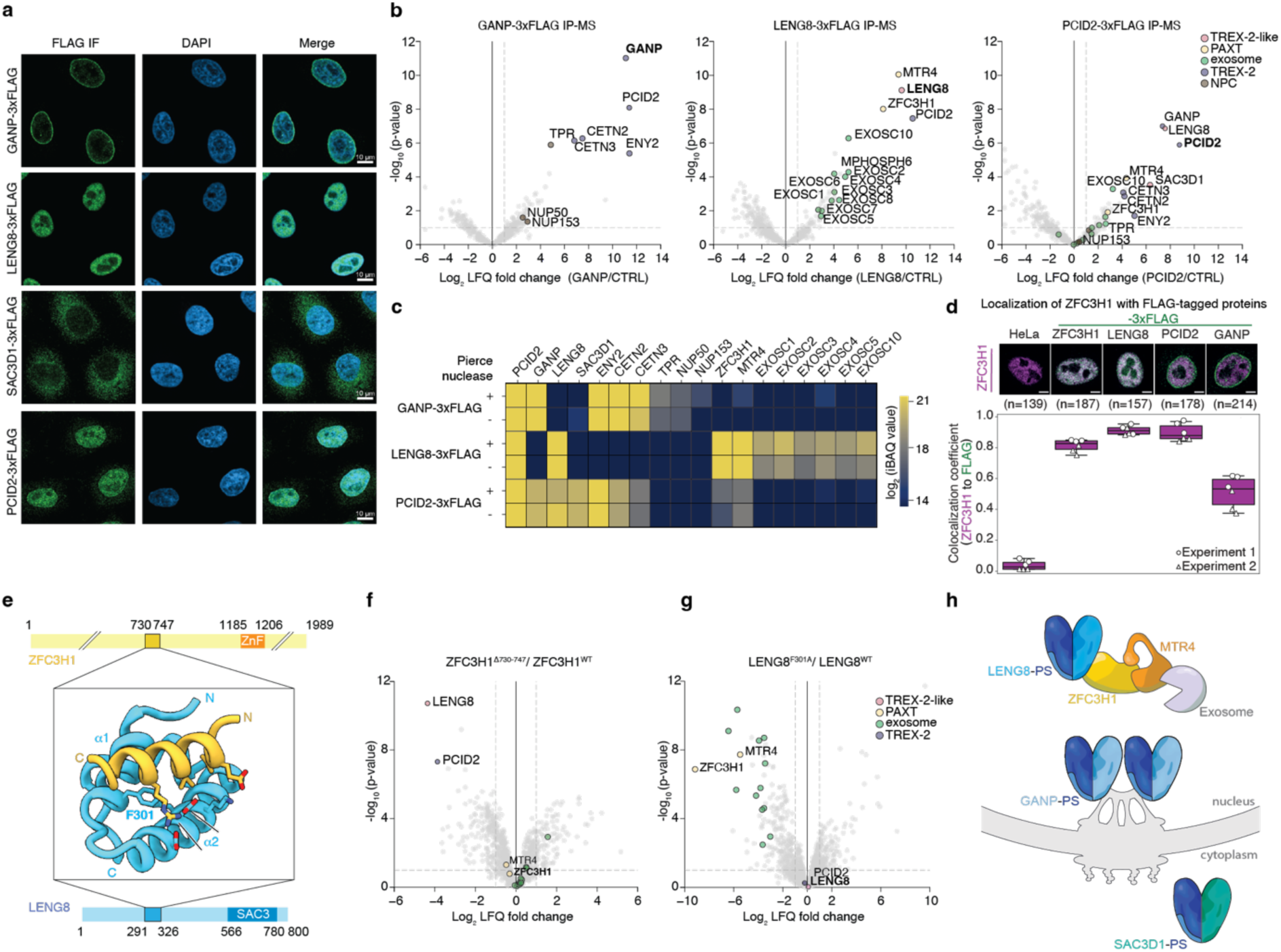
LENG8-PS constitutes a TREX-2-like module of PAXT. **a.** Immuno-localization analyses of central TREX-2 and TREX-2-like components. Anti-FLAG antibody (left column)- and DAPI (mid column)-stained HeLa cell lines, expressing C-terminally 3xFLAG tagged endogenous GANP (top), LENG8 (upper mid), SAC3D1 (lower mid) or PCID2 (bottom). Right column shows merges of FLAG and DAPI signals. **b.** Volcano plots of FLAG IP-MS analyses of extracts from 3xFLAG-tagged GANP (left), LENG8 (mid) or PCID2 (right) cells from (a). Log_2_ fold Label-Free Quantification (LFQ) changes of interactor signals in the individual IPs over their maternal HeLa cell line control (x-axes) were plotted against −log_10_ of Student’s t-test p-values calculated over triplicate data sets. TREX-2-like-, PAXT-, exosome-, TREX-2- and NPC-components are color-coded and labeled. For detection of SEM1 in the 3xFLAG-tagged LENG8 IP, chymotrypsin-treated IP-MS samples were employed (see Supplementary Fig. 4i). **c.** Heatmap of mean Intensity-Based Absolute Quantification (iBAQ) values from triplicate 3xFLAG-tagged GANP, LENG8, and PCID2 IP experiments, conducted without (-) or with (+) Pierce universal nuclease treatment. Displayed proteins as in (b). Note than only EXOSC1-EXOSC5, EXOSC10 subunits are shown. iBAQ values from control IP experiments were subtracted those of the experimental values. **d.** Median weighed co-localization coefficients between ZFC3H1- and FLAG-IF signals of 3xFLAG-tagged ZFC3H1, LENG8, PCID2 and GANP cell lines from Supplementary Fig. 3e. Maternal HeLa cells were used as a negative control. Red (re-colored in magenta) and green channels were used for the detection of ZFC3H1- and FLAG-signals, respectively. Each circle and triangle represent the colocalization coefficient calculated per image with three images from two independent experiment analyzed. Example cells with two color staining overlap and total numbers (n) of cells imaged are indicated above the plot. For examples of full co-staining see Supplementary Fig. 3e. **e.** AlphaFold2 model of interacting regions of ZFC3H1 (top) and LENG8 (bottom). The conserved phenylalanine at position 301 (F301) of LENG8, which is critical for interaction, is highlighted. Signature Zinc Finger (ZnF)- and Sac3- (SAC3) domains of ZFC3H1 and LENG8, respectively, are displayed. **f.** Volcano plot as in (b) but displaying log_2_ fold changes of mean LFQ protein intensities of ZFC3H^Δ730−747^-3xFLAG relative to ZFC3H1^WT^-3xFLAG sample data. Constructs were expressed in HeLa ZFC3H1-2xHA-dTAG cells treated with dTAG^V^-1 to deplete endogenous ZFC3H1. LENG8, PCID2 as well as PAXT- and exosome-components are color coded (see (**g**)). **g.** As in (**f**) but for LENG8^F301A^-3xFLAG relative to LENG8^WT^-3xFLAG sample data. Constructs were expressed in HeLa LENG8-2xHA-dTAG cells treated with dTAG^V^-1 to deplete endogenous LENG8. **h.** Cartoon depicting localization-distinct TREX-2-like modules. LENG8-PS interacts with PAXT in the nucleoplasm, GANP-PS associates with the NPC and SAC3D1-PS is cytoplasmic.

Owing to the critical role of UAP56 in pA^+^ RNP export via TREX-2, we hypothesized that LENG8 and SAC3D1 might target UAP56-bound RNPs to different cellular fates. To address this, we first explored the structure-function relationships of LENG8 or SAC3D1 with UAP56 *in vitro*. Indeed, as previously achieved for GANP^7^, we could purify stable recombinant complexes of SAC3 domain containing constructs of LENG8^491–800^ or SAC3D^148–404^ in the presence of PCID2–SEM1 (constituting LENG8-PS^M^ or SAC3D1-PS^M^), and to reconstitute interactions with UAP56 in the presence of the non-hydrolysable ATP analogue AMP-PNP and a 15-nucleotide poly-U RNA substrate (Supplementary Fig. 1b,c, lanes 1-5). Like the ability of TREX-2^M^ to stimulate the apparent ATPase rate of UAP56 in the presence of RNA and ATP^7^, *in vitro* ATPase assays revealed ∼290x and ∼60x stimulatory effects on the ATPase rate of UAP56 by LENG8–PS^M^ and SAC3D1–PS^M^, respectively (Fig. 1c,d; Supplementary Fig. 1d,e). Moreover, substituting a highly conserved arginine residue, in the LENG8 wedge loop, with an alanine (R563A) (Fig. 1b, Supplementary Fig. 1f)^7^, did not impact UAP56 binding (Supplementary Fig. 1g), yet largely abrogated the ATPase stimulatory activity of UAP56 (Fig. 1c). Finally, to test whether LENG8–PS^M^ and SAC3D1–PS^M^, like TREX-2^M^ (ref.^7^), would promote the release of RNA from UAP56, we incubated UAP56 with RNA and ATP to form UAP56–ADP–Pi–RNA complexes, which we immobilized on streptavidin beads via the biotinylated RNA^7^. These complexes were then challenged with either LENG8–PS^M^ or SAC3D1–PS^M^, revealing that both LENG8–PS^M^ or SAC3D1– PS^M^ discharged UAP56 efficiently (Fig. 1e). We conclude that TREX-2-like complexes, similar to TREX-2, bind UAP56 and trigger the release of its bound RNA through a conserved mechanism.

While previous structural studies of UAP56–TREX-2^M^ complexes had revealed their protein-protein interfaces, it remained unclear how the wedge loop functions in releasing UAP56 from RNA. To investigate the molecular basis for this, we analyzed LENG8–PS^M^ and SAC3D1–PS^M^ complexes with UAP56, in the presence of 15 U RNA and AMP-PNP, using cryo-EM. Unexpectedly, two-dimensional class averages of both complexes revealed UAP56 in a closed, RNA-bound state, prior to its release via the wedge loop, which was different to our previously reported UAP56–TREX-2^M^ structure, where UAP56 was captured after RNA release (Fig. 1f)^7^. While a severe bias in particle orientation prevented us from determining a three-dimensional structure of the UAP56–LENG8–PS^M^ complex, we successfully resolved the 2.6 Å cryo-EM structure of the UAP56–SAC3D1–PS^M^ complex in the pre-RNA release state (Fig. 1g, Supplementary Fig. 2, Supplementary table 1). This structure shared key architectural features with UAP56–TREX-2^M^, including the anchoring of UAP56’s N-terminal domain (NTD) at the base of the SAC3D1–PS complex. Truncating the UAP56 NTD reduced the affinity of UAP56 for both SAC3D1–PS^M^ and LENG8–PS^M^ by more than 30-fold, as addressed by Grating Coupled Interferometry and *in vitro* pulldown assays (Supplementary Fig. 1b,c, lanes 6-7; 1h). Hence, the UAP56 NTD is similarly important for TREX-2-like- and TREX-2-complex^7^ interactions. In addition, the UAP56–SAC3D1–PS^M^ structure provided insights into the action of the wedge loop. In the structure, this region (Fig. 1b, Supplementary Fig. 1f, residues Y100-P111) is bound near the two RecA lobes through largely electrostatic interactions between the peptide backbone and UAP56 residues R135 in RecA1 and K334 in RecA2 (Fig. 1h). Here, the critical R102 wedge loop residue in SAC3D1 forms a hydrogen bond with UAP56 E354, which pre-positions R102 adjacent to F381 in the UAP56 RecA2 lobe, priming it to displace this residue in a next step to release UAP56 from RNA (Fig. 1h,i and ref.^7^).

We conclude that human cells, besides TREX-2, contain two structurally and biochemically equivalent complexes, which can bind UAP56 and through the conserved wedge loops of their respective SAC3 domains dislodge UAP56 from RNA.

### LENG8-PS defines a distinct physical module of PAXT

Our biochemical and structural analyses suggested that the GANP-PS, LENG8-PS and SAC3D1-PS complexes can individually act on UAP56. To address where these complexes act *in vivo*, we generated HeLa cell lines^22^, stably expressing C-terminally 3xFLAG-tagged versions of endogenous GANP, LENG8, or SAC3D1 as well as the common subunit PCID2 (Supplementary Fig. 3a, lanes 3-6), and subjected these to immunofluorescence (IF) microscopy using a FLAG-antibody. As previously reported, GANP was found primarily at the nuclear envelope, consistent with its NPC association^6,23^ (Fig. 2a, top panel). In contrast, LENG8 and SAC3D1 localized to the nucleoplasm and the cytoplasm, respectively (Fig. 2a, mid panels). In agreement with its presumed presence in all three complexes, PCID2 distributed between the nucleoplasm, the nuclear envelope and the cytoplasm (Fig. 2a, bottom panel).

With our focus on nuclear RNA sorting, we scrutinized the TREX-2 and LENG8-PS complexes in more detail, performing immunoprecipitation/mass spectrometry (IP-MS) analyses of the GANP-, LENG8- and PCID2-3xFLAG proteins. Stringent IP conditions were used to enrich for high-affinity interactors. In the GANP-3xFLAG IP, this yielded high amounts of PCID2, in addition to the ENY2 and CETN2/3 subunits of TREX-2 (ref^5^), and the nuclear pore basket protein TPR^23^ (Fig. 2b, left panel). While LENG8-3xFLAG also precipitated PCID2, this IP was instead enriched for the PAXT core factors ZFC3H1 and MTR4 along with nuclear exosome subunits (Fig. 2b, mid panel, note that SEM1 was also detected applying an alternative protocol (Supplementary Fig. 4i)). Finally, reflecting on its dual interaction with GANP and LENG8, PCID2-3xFLAG precipitated these proteins and their respective interactors (Fig. 2b, right panel). Subjecting our IP-MS experiments to RNase-treatment revealed that the mentioned interactions were not facilitated by RNA (Fig. 2c, S3b). Moreover, analyzing mean enrichments across IP experiments displayed near-identical interaction levels of GANP with TREX-2 components, while LENG8 precipitated similar amounts of PCID2 as well as the core PAXT factors ZFC3H1 and MTR4 (Fig. 2c, Supplementary Fig. 3a, lanes 7-12; Supplementary table 2). This indicated that LENG8-PS might be a stable module of PAXT. To address this notion further, we generated ZFC3H1-3xFLAG cells (Supplementary Fig. 3a, lane 2) and conducted IP-MS, which revealed abundant and RNase-resistant amounts of LENG8 and PCID2 (Supplementary Fig. 3a,c,d). Calculating mean enrichments of proteins showed that, along with exosome subunits, LENG8 and PCID2 were sub-abundant to MTR4 (Supplementary Fig. 3d), which was in agreement with the reported presence of inactive nuclear ZFC3H1-MTR4 dimers^12^. Finally, association of LENG8 with ZFC3H1 was mirrored by their measured nuclear co-localization, as revealed by immunostaining of LENG8-3xFLAG cells using FLAG- and ZFC3H1-specific antibodies. Here, LENG8 and PCID2, but not GANP, displayed weighed co-localization coefficients with ZFC3H1 in the nucleoplasm of over 90% (Fig. 2d, Supplementary Fig. 3e, Supplementary table 2).

Having established a physical link between LENG8 and ZFC3H1, we inquired whether LENG8 interacts with MTR4 and the exosome via ZFC3H1. Indeed, rapid ZFC3H1 depletion, using the FKBP12^F36V^-degron (dTAG) ^24^ (Supplementary Fig. 4a, left panel), prevented interactions of MTR4 and the exosome component EXOSC10 with LENG8-3xFLAG (Supplementary Fig. 4a, right panel). To identify a possible ZFC3H1 interaction site on LENG8, we employed AlphaFold2, which revealed a conserved motif of two α-helices (residues 288-342) in the otherwise unstructured region N-terminal of the SAC3 domain (Supplementary Fig. 4b,c). A direct interaction was predicted to take place between this helical region of LENG8 and a single α-helix (residues 730-747) of ZFC3H1 (Fig 2e, Supplementary Fig. 4d). Validating this prediction, IP-MS analysis of a ZFC3H1^Δ730–747^– compared to a ZFC3H1^WT^-3xFLAG construct, both expressed upon depletion of endogenous ZFC3H1, demonstrated the specific loss of LENG8 and PCID2 (Fig. 2f, Supplementary Fig. 4e). Likewise, mutation of the central LENG8 phenylalanine (F301) at the LENG8–ZFC3H1 interface (Fig. 2e) to an alanine lead to the loss of ZFC3H1, MTR4 and exosome subunits, in a comparative IP-MS analysis of LENG8^WT^ versus LENG8^F301A^-3xFLAG constructs (Fig. 2g, Supplementary Fig. 4f-h, Supplementary table 2).

Taken together, we conclude that the TREX-2-like LENG8-PS complex constitutes a physical module of the nucleoplasmic PAXT connection (Fig. 2h).

### LENG8-PS functions in PAXT-mediated control of nuclear pA^+^ RNAs

Given the physical link between ZFC3H1 and LENG8, we next sought to probe its functional relevance. Expression of ZFC3H1WT-3xFLAG, but not its ZFC3H1^Δ730–747^ LENG8 binding-deficient variant (Supplementary Fig. 5a), suppressed select PAXT substrates from accumulating after rapid depletion of endogenous ZFC3H1 (Fig. 3a). Likewise, PAXT substrate accumulation, following rapid depletion of endogenous LENG8, was suppressed by expression of LENG8^WT^-3xFLAG but not by the ZFC3H1 binding mutant LENG8^F301A^ (Fig. 3b, Supplementary Fig. 5b).

**Figure 3.**
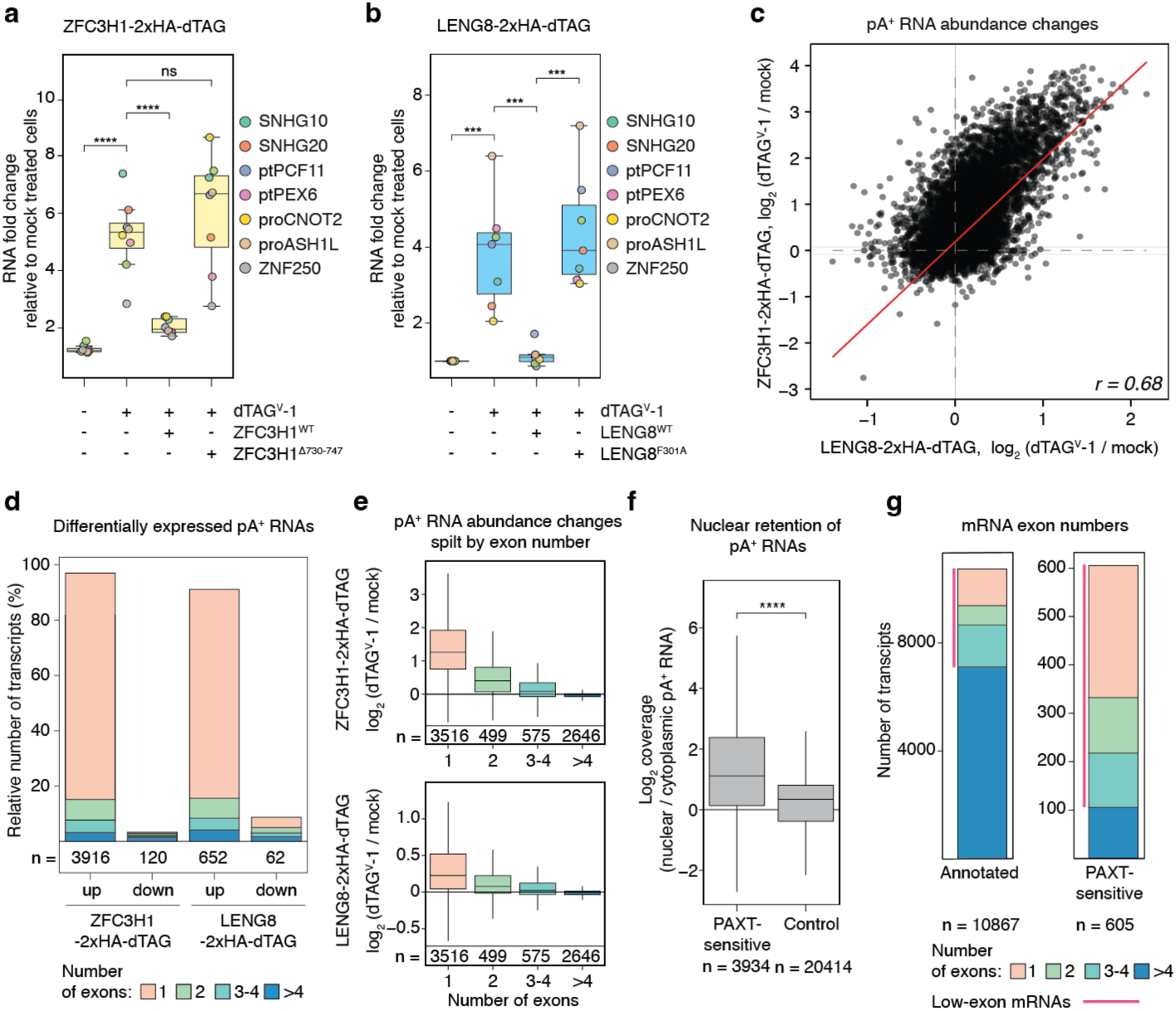
LENG8-PS is a functional module of PAXT. **a.** Boxplots of RT-qPCR analysis of select PAXT substrates from ZFC3H1-2xHA-dTAG cells treated with dTAG^V^-1 for 4 hours (+), or not (-), and complemented by ZFC3H1^WT^ or ZFC3H1^Δ730–747^ expression as indicated. RT-qPCR values were plotted relative to mock-treated HeLa cells and normalized to GAPDH RNA levels. Colored dots denote the mean value of three biological replicates. Student’s two-sided t-tests were calculated between conditions, combining the results for all RNAs tested (***p-value<0.001, ****p-value<0.0001). Similar statistical tests were conducted for the subsequent qRT-PCR analyses and with p-values represented in the same manner. The suffix ‘pt’ indicates ‘prematurely terminated’. **b.** As (a) but for LENG8-2xHA-dTAG cells treated with dTAG^V^-1 for 24 hours (+), or not (-), and complemented by LENG8^WT^ or LENG8^F301A^ expression as indicated. **c.** Scatter plot of pA^+^ RNA log_2_ fold abundance changes resulting from treatment of ZFC3H1- (y-axis) or LENG8-2xHA-dTAG (x-axis) cells with 500 nM of dTAG^V^-1 for 4 hours. Trendline is shown in red and Pearson correlation coefficient (*r*) is plotted in the low right corner. **d.** Counts of differentially expressed pA^+^ RNAs after treatment of ZFC3H1- or LENG8-2xHA-dTAG cells with dTAG^V^-1 for 4 hours, Transcripts are color-coded by their numbers of exons. DESeq2 (−log_10_ (adj. p-value) > 1) was employed and RNAs with log_2_ (fold change) > 0.5 and < −0.5 were counted as upregulated and downregulated, respectively, both here and in subsequent analyses. **e.** Boxplots of pA^+^ RNA log_2_ fold abundance changes after dTAG^V^-1 treatment for 4 hours of ZFC3H1-2xHA-dTAG (top) or LENG8-2xHA-dTAG (bottom) cells. All RNAs with measurable fold changes (−log_10_ (adj. p-value) > 1) in either of the depletions were included and grouped by their contained exon numbers. **f.** Boxplots of log_2_-scaled nuclear/cytoplasmic ratios in unperturbed HeLa cells of upregulated (log_2_ (fold change) > 0.5, −log_10_ (adj. p-value) > 1) ZFC3H1- or LENG8-sensitive pA^+^ RNAs compared to all other annotated RNAs. Welch’s t-test were calculated between conditions, combining the results for all RNAs tested (**** p-value <0.0001). **g.** Counts of all annotated (left) or PAXT-sensitive (right) mRNAs stratified by their contained exon numbers. PAXT-sensitivity was scored as increased transcript levels upon ZFC3H1- or LENG8-depletion. mRNAs containing 1 to 4 exons (classified as low-exon) are marked with a pink bar.

Since these analyses corroborated a role for LENG8 in PAXT function, we obtained a transcriptome-wide view of this relationship by sequencing pA^+^ RNA from cells rapidly depleted for either LENG8 or ZFC3H1 (Supplementary Fig. 5c-d). PAXT substrates include numerous prematurely terminated transcripts (PTTs), deriving from transcription start site (TSS)-proximal regions of protein-coding genes^9,13^. To incorporate these in our analysis, we identified TUs displaying such increased pA^+^ RNA coverage and intersected data with pA^+^ RNA 3’end peaks previously identified upon depletion of ZFC3H1 (Supplementary Fig. 5e; ref.^25^; see also Methods). As a result, 1202 PTTs were defined (see Supplementary Fig. 5f for an example) and included in our HeLa transcriptome annotation^26^ (Supplementary Fig. 5g). Although numbers and levels of affected RNAs were higher for the ZFC3H1 depletion, individual LENG8- and ZFC3H1-sensitive transcripts were strongly correlated (Fig. 3c), consistent with a shared pathway. This was substantiated by sequencing pA^+^ RNA from ZFC3H1-depleted cells, exogenously expressing either ZFC3H1WT or the LENG8-binding mutant ZFC3H1^Δ730–747^ (Supplementary Fig. 5h); only ZFC3H1^WT^ suppressed the accumulation of PAXT-substrates (Supplementary Fig. 5i).

Differential expression (using DESeq2^27^) analysis of ZFC3H1 and LENG8 rapid depletion data yielded, in agreement with previous PAXT studies^9,12,14^, upregulated promoter upstream transcripts (PROMPTs)^28^, PTTs, other non-coding (nc) RNAs and a minor fraction of full-length mRNAs (Supplementary Fig. 5j; Supplementary table 3). In line with this, the majority of ZFC3H1- and LENG8-substrates contained only one or a few exons (Fig. 3d) and for both depletion conditions RNA accumulation levels decreased with increasing exon number (Fig. 3e) and mature RNA length (Supplementary Fig. 5k). In addition to decay, ZFC3H1 also contributes to nuclear retention of pA^+^ RNA^12,13,15^. To analyze the impact of ZFC3H1 or LENG8 on transcript fate, we therefore performed nuclear/cytoplasmic pA^+^ RNAseq obtained by fractionating cells before or after the rapid depletion of these factors (Supplementary Fig. 6a-b). Consistent with the RNA-retention capacity of ZFC3H1, PAXT-sensitive RNAs displayed higher nuclear-to-cytoplasmic ratios in unperturbed HeLa cells than the remaining pA^+^ transcriptome (Fig. 3f). Clustering all transcripts by their depletion-dependent nuclear or cytoplasmic content changes demonstrated that more than half of the displayed RNAs were immune to ZFC3H1 or LENG8 depletion (Supplementary Fig. 6c, left panel, Cluster 1, Supplemetary table 3; Methods). While most of these transcripts were accounted for by spliced mRNAs with multiple exons (Supplementary Fig. 6c, see mid panel for biotypes), a closer examination of the low-exon count RNAs from Cluster 1 revealed mild, but detectable, sensitivity to ZFC3H1 depletion (Supplementary Fig. 6d). Outside of Cluster 1, the remaining transcripts were, to variable extents, upregulated in both compartments upon ZFC3H1 and LENG8 depletions, demonstrating inefficient nuclear retention (Supplementary Fig. 6a, lanes 1-4 and 7-10; Supplementary Fig. 6c, left panel, Clusters 2-4). It therefore appears, that PAXT retains and mediates decay of short pA^+^ RNAs with few exons whereas transcripts escaping these fates are generally longer and more exon-rich.

Although the protein-coding fraction of analyzed transcripts was largely insensitive to PAXT, a minor subset of sensitive mRNAs was still detectable (Supplementary Fig. 6c, note biotypes of Clusters 2-4). Like non-coding PAXT substrates, these were primarily low-exon content (1-4 exons) transcripts (Fig. 3g), enriched in nuclei of unperturbed cells (Supplementary Fig. 6e). However, in the same condition, these short mRNAs were present at higher levels than their ncRNA counterparts (Supplementary Fig. 6f), suggesting that they, at least to some extent, have acquired means to fend off nuclear turnover (see Discussion). DESeq2 also identified 105 longer, multi-exonic mRNA outliers (>4 exons), that were PAXT-sensitive despite their larger exon number and length. As general PAXT targets (Fig. 3f), these transcripts showed increased nuclear-to-cytoplasmic ratios in unperturbed cells as compared to PAXT-insensitive controls (Supplementary Fig. 6g), suggesting that prolonged nuclear residence time may drive their sensitivity to PAXT. Notably, the LENG8 mRNA belongs to this transcript category, implying autoregulation of the PAXT pathway (see below). Since introns contribute to nuclear RNA retention^29–31^, we assessed PAXT-sensitive multi-exonic mRNAs (>4 exons) for reads spanning both 5′ and 3′ splice junctions (see Methods). Indeed, when compared to a control population, these transcripts were enriched for retained introns, including the previously described detained introns^32^ (Supplementary Fig. 6h). Moreover, upon ZFC3H1- or LENG8-depletion, PAXT-sensitive multi-exonic transcripts accumulated largely as incompletely spliced precursors in the nucleus and as fully spliced mRNAs in the cytoplasm (Supplementary Fig. 6i, j). Thus, upon PAXT impairment, this group of transcripts can be post-transcriptionally spliced and exported. Finally, we also identified an additional 3,209 PAXT-sensitive introns, including 161 detained introns (Supplementary table 4), for which the corresponding spliced mRNAs were PAXT-insensitive as measured by DESeq2. We therefore propose that, in addition to short RNAs, PAXT can also target multi-exonic transcripts, that are retained in the nucleus due to incomplete splicing. In certain cases, this alters the cellular levels of the mature transcript, the extent to which might differ between cell types.

Taking our cellular data together, we conclude that LENG8-PS constitutes a functional module of PAXT, which primarily targets nuclear pA^+^ RNAs with no, or only a few, introns as well as a minor pool of longer intron-containing transcripts. This overall retention/decay regime may be exploited for the regulation of select mRNAs.

### PAXT and TREX-2 contact UAP56-bound RNP to govern nuclear pA+ RNA fate

Having defined LENG8-PS as a central physical and functional module of PAXT, we investigated how LENG8’s ability to release UAP56 from RNA *in vitro* (Fig. 1e) affects its function *in vivo*. Indeed, the LENG8^R563A^ wedge loop mutant (Supplementary Fig. 5b), defective in releasing UAP56 from RNA *in vitro* (Fig. 1c, d), was impaired in binding RNA *in vivo* (Supplementary Fig. 7a) and in suppressing the upregulation of select PAXT targets sensitive to endogenous LENG8 depletion (Fig. 4a). Since the LENG8^R563A^-3xFLAG construct retained interactions with PAXT components (Supplementary Fig. 7b), we conclude that release of RNA from UAP56 is central for its PAXT-mediated turnover. Unexpectedly, however, the LENG8^H712F^ point mutant (Supplementary Fig. 5b, 7c), abolishing the interaction with PCID2 (Supplementary Fig. 7d-e, Supplementary table 1), partially rescued the LENG8 depletion phenotype (Fig. 4a), prompting us to re-evaluate the role of the PCID2-SEM1 module for LENG8 function in releasing UAP56-bound RNA. Curiously, the protein-protein interfaces between UAP56 and PCID2 in both TREX-2^M^ (ref.^7^) and the TREX-2^M^-like complexes involve only few specific interactions (Fig. 1f, g). We therefore compared the ATPase stimulatory activity of a SAC3-domain-containing protein, LENG8, itself, with that of the LENG8-PS^M^ complex. Surprisingly, LENG8 alone robustly stimulated the ATPase activity of both UAP56 and its close paralog EIF4A3 (Supplementary Fig. 7f, g). In contrast, LENG8–PS^M^ exhibited no effect on EIF4A3. Super positioning of the RNA-bound EIF4A3 onto the UAP56–SAC3D1–PS^M^ structure revealed clashes between EIF4A3 and PCID2, while this was not observed with the SAC3-domain or the wedge loop (Supplementary Fig. 7h). Thus, the PS module of TREX-2 and TREX-2–like complexes appears to ensure specificity for UAP56, presumably preventing other DExD-box proteins from being affected.

**Figure 4.**
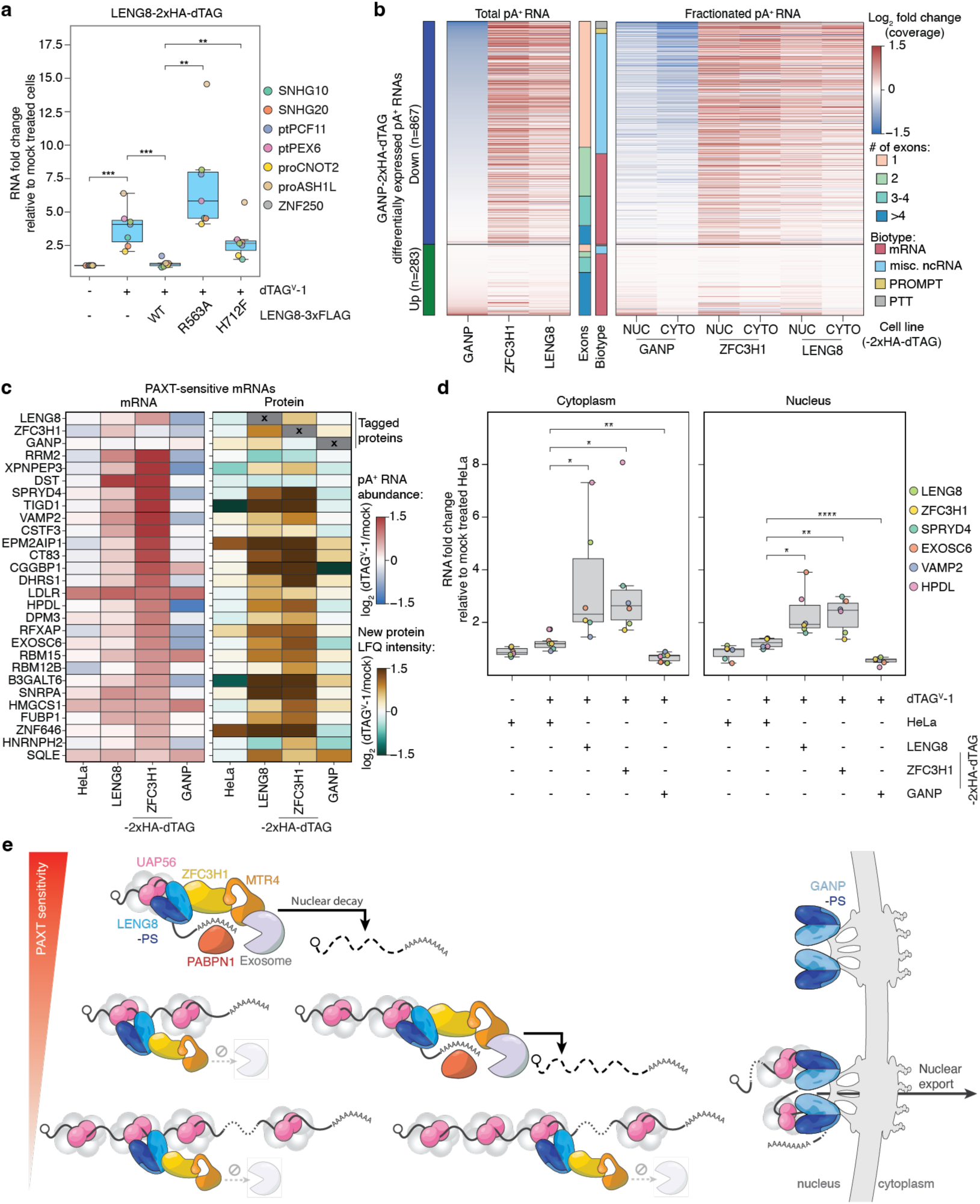
PAXT and TREX-2 compete for pA^+^ RNAs. **a.** Boxplots of RT-qPCR analysis of select PAXT substrates as in Fig. 3b, but employing LENG8^WT^, LENG8^R563A^ or LENG8^H712F^ constructs as indicated. **b.** Heatmaps as in Supplementary Fig. 6d, but of pA^+^ RNA log_2_ fold abundance changes in total (left panel) or nuclear (NUC) and cytoplasmic (CYTO) fractions (right panel) of samples of dTAG^V^-1-treated (4 hours) GANP-, ZFC3H1- or LENG8-2xHA-dTAG cells relative to maternal HeLa cells. Transcripts were clustered and ranked based on differential expression analysis of total GANP depletion sample. **c.** Heatmaps of log_2_ fold abundance changes of PAXT-sensitive mRNAs (DESeq2, see Fig. 3) (left) and their SILAC LFQ intensity changes (right) upon dTAG^V^-1 treatment of LENG8-, ZFC3H1- or GANP-2xHA-dTAG cells. mRNA–protein pairs, that were significantly differentially expressed (DESeq2 for mRNAs and DEP for proteins) with –log₁₀(p-value) > 1, are shown. Maternal HeLa cells are shown as controls. Heatmaps were sorted by descending mRNA sensitivity to ZFC3H1 depletion. LENG8-, ZFC3H1- and GANP-depletion induced mRNA and -SILAC abundance changes are displayed in the upper three rows. **d.** Boxplots of RT-qPCR analysis of select short TREX-2/PAXT substrates as well as the LENG8 and ZFC3H1 mRNAs in nuclear and cytoplasmic fractions of biochemically fractionated maternal HeLa cells or 2xHA-dTAG-tagged cell lines subjected to dTAG^V^-1-induced depletion of LENG8, ZFC3H1 or GANP for 4 hours. RNA fold changes were calculated relative to levels of the non-treated HeLa control and normalized to levels of GAPDH mRNA, separately for cytoplasmic (left) and nuclear (right) fractions. **e.** Model of nuclear pA^+^ RNA sorting. See text for details.

Taken together, the above experiments demonstrated the targeting of UAP56-bound RNPs by PAXT-LENG8-PS *in vivo*, suggesting that UAP56, while being an RNA export factor, is also crucial for nuclear RNA decay. To interrogate possible relations between these two fates of UAP56-bound RNPs, we first constructed GANP-2xHA-dTAG cells, enabling the rapid reduction of GANP protein (Supplementary Fig. 5c, Methods), and analyzed its consequence by pA^+^ RNAseq (Supplementary Fig. 5d). In agreement with previous studies^23,33^, this mainly resulted in the downregulation of short pA^+^ RNAs with few introns (Fig. 4b, Supplementary Fig. 7i-j), resembling PAXT-sensitive transcripts (Fig. 3d; Supplementary Fig. 6a). Indeed, GANP-sensitive RNAs were largely upregulated upon depletion of ZFC3H1 or LENG8 (Fig. 4b, Supplementary Fig. 7k). We therefore propose that the suppressive effect of GANP reduction reflects PAXT-mediated degradation of export-restricted transcripts. Consistently, our fractionated pA^+^ RNAseq data displayed the coinciding cytoplasmic accumulation of these RNAs following LENG8 or ZFC3H1 depletion (Fig. 4b, right panel). In contrast, GANP reduction did not significantly impact levels of nuclear retained multi-exonic (>4 exons) PAXT substrates (Supplementary Fig. 7l; notice the LENG8 mRNA defying this trend, see also below).

To further elaborate on an apparent competition of TREX-2 and PAXT for their target transcripts, we monitored protein production in TREX-2- or PAXT-perturbed cells. As expected, GANP-depletion impaired global protein synthesis as revealed by decreased puromycin incorporation (Supplementary Fig. 8a). Less intuitive, but consistent with prior reports on ZFC3H1(ref.^12,25^), both LENG8- and ZFC3H1-depletions also decreased new protein synthesis, possibly due to the overloading of ribosomes by ncRNAs escaping from the nucleus. We characterized these translational alterations further through the quantitative analysis of nascent protein synthesis using Stable Isotope Labelling by Amino acids in Cell culture (SILAC) followed by MS^34^ (Supplementary table 5). Although only a subset of proteins matching PAXT-sensitive mRNAs were detectable, LENG8- or ZFC3H1-depletions generally increased their de novo peptide synthesis, whereas GANP depletion had little or opposing effects (Fig. 4c; Supplementary Fig. 8b). We note that this occurred despite the general conditions of decreased protein synthesis (Supplementary Fig. 8a). Proteins with increased synthesis upon LENG8- or ZFC3H1-depletions were enriched for ‘RNA binding proteins’ and ‘RNA processing regulators’ (Supplementary table 5), including increased LENG8 protein synthesis upon ZFC3H1 depletion and increased ZFC3H1 protein synthesis in case of LENG8 depletion (Fig.4c, top rows). As the ZFC3H1 mRNA did not pass the DESeq2 analysis threshold (Supplementary table 3), we employed RT-qPCR to reveal that rapid depletion of LENG8 or ZFC3H1 led to the nuclear and cytoplasmic accumulation of their own mRNAs, whereas GANP depletion reduced such levels (Fig. 4d). This supports the previous notion, that retained mRNAs, as the LENG8 mRNA, are subject to regulation by the PAXT pathway itself.

In conclusion, two structurally and functionally similar modules, LENG8-PS and GANP-PS, are critical interpreters of UAP56-bound nuclear pA^+^ RNPs. Nucleoplasmic PAXT and nuclear pore-associated TREX-2 utilize these biochemically redundant modules to control nuclear pA^+^ RNA homeostasis by facilitating decay or export, respectively.

## DISCUSSION

In this study, we demonstrate that short and low exon-content pA^+^ RNAs, hosted in UAP56-bound RNPs, are highly susceptible to PAXT-mediated nuclear turnover. Instead, TREX-2 seemingly grants export to any pA^+^ RNA provided that this pA^+^ RNP can make its way to the NPC. Based on these findings, we propose a general model for nuclear pA^+^ RNA fate determination (Fig. 4e). Both TREX-2 and PAXT engage UAP56-bound pA^+^ RNPs indiscriminately but due to their nucleoplasmic localization the majority of pA^+^ RNPs would first encounter PAXT. However, being an adaptor of the 3’-5’ exonucleolytic exosome, PAXT targeting only translates into efficient decay if it occurs in the vicinity of the PABPN1-bound pA^+^ RNA 3’end. This condition greatly sensitizes short transcripts. In contrast, for longer RNAs, compacted into larger RNPs with multiple UAP56s, a PAXT encounter, while resulting in the release of a UAP56 molecule, may not translate into RNA decay, in turn increasing the probability of nuclear export via TREX-2. In support of this model, a recent study found that the widespread interaction of ZFC3H1 with long and multiply spliced mRNAs did not affect transcript levels upon ZFC3H1 depletion^15^. That said, targeting of LENG8-PS to decay-insensitive transcripts might nevertheless play a role in maintaining sufficient levels of free nuclear UAP56, without which mRNA biogenesis defects, R-loop formation and genomic instability would prevail^35,36^. The importance of such UAP56 recycling might in fact explain why budding yeast, where the PAXT complex has been lost, still harbors the LENG8-PS homolog^21^.

Although general, the proposed model may be bypassed by specific transcripts. For example, short functional pA^+^ mRNAs, including stress-induced transcripts, must overcome nuclear decay. How this is achieved remains an open area of research. Perhaps stress-induced mRNP reorganization offers protection to the RNA 3′end, or perhaps their robust transcriptional upregulation ensures that, despite ample nucleoplasmic decay, a sufficient number of RNPs still reaches the NPC. Consistent with the latter notion, we find that PAXT-sensitive short mRNAs are expressed at higher levels than their ncRNA counterparts (Supplementary Fig. 6f). Finally, decay might also be short-circuited by gene gating, positioning a given locus in proximity to the NPC, as was recently demonstrated for c-MYC^37,38^.

We also find a small subset of multi-exonic transcripts, that are sensitive to PAXT-mediated decay, violating the general nuclear pA^+^ RNA decay regime (Fig. 3g). We suggest that this sensitivity is enhanced by prolonged residence in the nucleus (Supplementary Fig. 6e, g), exacerbated by intron presence (Supplementary Fig. 6h). In line with nuclear retention of (pre-)mRNA being able to play a regulatory role^29–31^, our data provide evidence that the PAXT system itself is subject to such control (Fig. 4c). Taken together, the PAXT axis, which primarily targets short, non-functional pA^+^ RNAs, has been co-opted to regulate a subset of mRNAs that are sensitized either by low exon content or prolonged nuclear retention.

In closing, we propose that the major opposing fates of nuclear pA^+^ RNA - export and decay - exploit a shared molecular logic. At its center is RNA-bound UAP56 and its highly regulated release from the pA^+^ RNP. Our findings blur previously held categorizations of proteins being specific RNA export- or decay-factors and highlight the fierce competition for common pA^+^ RNPs features that together ensures faithful gene expression.

## ACKNOWLEDGEMENTS

Members of the T.H.J., C.P., and J.B. laboratories are thanked for fruitful discussions. Anne Aagaard and Dorthe Riishøj are thanked for expert technical assistance and Will Garland for critical reading of the manuscript. Staff at the Protein Technologies Facility at the Vienna BioCenter Core Facilities (VBCF) is thanked for assistance with protein production. Staff at the VBCF Electron Microscopy Facility, particularly T. Heuser and H. Kotisch, are thanked for support, data collection and maintaining facilities. Staff at the VBC Molecular Biology Service and the CLIP cluster (https://clip.science) are thanked for reagents and computation support, respectively. Work in the T.H.J. laboratory was supported by the Danish National Research Council, the Lundbeck Foundation (R1982015-172) and the Novo Nordisk Foundation (NNF18OC0033380 and ExoAdapt grant 31199). Research in the J.B. laboratory was supported by the Austrian Academy of Sciences and Austrian Science Fund (W1207). Research in the laboratory of C.P. was supported by Boehringer Ingelheim, the European Research Council under the Horizon 2020 research and innovation programme (ERC-2020-STG 949081 RNApaxport) and by the Austrian Science Fund (FWF) doc.funds program DOC177-B (RNA@core: Molecular mechanisms in RNA biology). A.B. was supported by the Marie Sklodowska-Curie Individual Fellowship (EXOonRNA, 101026781) and a Lundbeck Foundation Experiment Grant (R346-2020-1610). U.H. was supported by a Marie Sklodowska-Curie fellowship (896416) and an EMBO long-term fellowship (ALTF_1175-2019). A.L. and M.G. were supported by Boehringer-Ingelheim Fonds fellowships. For the purpose of open access, the authors have applied for a CC BY public copyright license to any author accepted manuscript version arising from this publication. The funders had no role in study design, data collection and analysis, decision to publish or preparation of the manuscript.

## AUTHOR CONTRIBUTIONS

T.H.J., C.P., U.H., J.B., A.L., M.G. and A.B. conceptualized the study; M.G., U.H. and L.F. performed *in vitro* experiments; U.H., M.G., C.P. collected and analyzed cryo-EM data; A.B., A.L. and Y.D. performed immunoprecipitations; A.L performed immunofluorescence imaging and data analysis; A.B., L.J., and J.S.A performed whole cell SILAC-MS and data analysis; L.J., and J.S. performed MS; A.B. and A.L. collected RNA samples for sequencing; A.B., A.L., D.J., P.P. and L.T. generated cell lines; J.O.R., A.B. and A.L. performed computational analyses of sequencing and MS data; A.B., U.H., A.L., M.G., C.P., J.B., and T.H.J wrote the manuscript with input from all co-authors; T.H.J., C.P., J.B. and A.B. acquired funding; T.H.J., C.P. and J.B. supervised the study.

## METHODS

### DNA sequences

All oligonucleotides plasmid vectors are annotated in Supplementary table 6.

### Protein purification

*<u>UAP56 and UAP56^ΔNTD^:</u>* His-tagged UAP56 constructs (10xHis-3C-UAP56 or 10xHis-3C-UAP56ΔNTD, residues 44–428) were expressed in *E. coli* BL21 DE3 RIL using autoinduction media at 37°C for 16 hours. Following harvest, cells were resuspended in lysis buffer (25 mM HEPES pH 7.9, 5% glycerol, 300 mM NaCl, 20 mM imidazole, 0.05% Tween-20, and protease inhibitors), disrupted via sonication, and clarified by centrifugation. The supernatant was sequentially filtered through 1 µm and 0.45 µm filters before affinity purification on a HisTrap HP 5 mL column (Cytiva), equilibrated in buffer A (25 mM HEPES pH 7.9, 5% glycerol, 300 mM NaCl, 20 mM imidazole). After washing with buffer A supplemented with 70 mM imidazole, bound proteins were eluted using a linear gradient of imidazole (70–200 mM in buffer A). Peak fractions were diluted in buffer B (25 mM HEPES pH 7.9, 5% glycerol, 1 mM DTT) to reduce the NaCl concentration to 100 mM and subsequently subjected to anion-exchange chromatography on a HiTrapQ 5 mL column (Cytiva), pre-equilibrated with buffer B. Elution was performed with a linear NaCl gradient (100–500 mM). Fractions containing UAP56 were concentrated and further purified via size-exclusion chromatography using a HiLoad 16/600 Superdex 200 pg column (Cytiva), equilibrated in buffer C (25 mM HEPES pH 7.9, 5% glycerol, 100 mM NaCl, 1 mM DTT). Peak fractions containing the purified protein were pooled, concentrated, flash-frozen, and stored at −80°C.

*<u>LENG8–PS^M^ and SAC3D1–PS^M^:</u>* Expression constructs encoding LENG8–PS^M^ (10His-MBP-LENG8^491–800^, 3xV5-PCID2, SEM1), SAC3D1–PS^M^ (10His-MBP-SAC3D1^48–404^, 3xV5-PCID2, SEM1), SAC3D1–PCID2-UAP56-UCM-N-UBM–SEM1 (10His-MBP-SAC3D1^48–404^, 3xV5-PCID2-UAP56-UCM-N-UBM, SEM1), LENG8-PCID2-UAP56-N-UBM_SEM1 (10His-MBP-LENG8^491–800^, 3xV5-PCID2-UAP56-N-UBM, SEM1) and 10His-MBP-LENG8^501–800^ mutants were introduced into *E. coli* BL21 DE3 RIL. Cultures were grown in LB medium at 37°C to OD600 ∼1.0, at which point expression was induced with 0.5 mM IPTG, followed by overnight incubation at 18°C. Cells were collected, lysed by sonication, and clarified by centrifugation. The supernatant was filtered (1 µm and 0.45 µm) and loaded onto a HisTrap HP 5 mL column equilibrated with buffer A, followed by washing and elution using a linear imidazole gradient up to 300 mM. Peak fractions were diluted to 50 mM NaCl in buffer B and subjected to anion-exchange purification on a HiTrapQ HP 5 mL column. After washing, complexes were eluted with a NaCl gradient (100–500 mM). Size-exclusion chromatography using a HiLoad 16/600 Superdex 200 pg column (Cytiva) in buffer C containing 250 mM NaCl yielded the final purified complex, which was concentrated, flash-frozen, and stored at −80°C.

*EIF4A3:* Recombinant eIF4A3 was purified as described previously^1^.

### Pull down experiments using recombinant proteins

*<u>UAP56–LENG8 and UAP56–SAC3D1-PS^M^ pulldown:</u>* MBP-tagged LENG8-PS^M^ or SAC3D1-PS^M^ was incubated with a 4-fold molar excess of UAP56 or UAP56^ΔNTD^ in buffer D (25 mM HEPES pH 7.9, 40 mM NaCl, 5% glycerol, 0.01% IGEPAL CA630, 1 mM MgCl2, 1 mM TCEP), with or without 50 μM 15U RNA and 1 mM AMP-PNP. Reactions were mixed by rotation at 4°C for 1 hour before adding 30 μL of pre-equilibrated amylose resin (E8021S, NEB). After an additional 1-hour incubation at 4°C, unbound proteins were removed by centrifugation (1500 × g, 2 min, 4°C) and three washes with buffer D. Bound proteins were eluted by incubation at 4 °C for 1 hour in buffer D supplemented with 100 mM maltose. Input and elution fractions were analyzed via SDS-PAGE (4–12% gradient) and visualized by Coomassie staining.

*<u>RNA unclamping assay:</u>* Biotinylated 15U RNA (33 µM) was mixed with recombinant UAP56 (10 µM) and 1 mM ATP in buffer E (20 mM HEPES pH 7.9, 40 mM KCl, 2 mM MgCl2, 5% glycerol, 0.1% IGEPAL CA630). This mixture was incubated with 20 µL NeutrAvidin™ Agarose beads (#29202, ThermoScientific), pre-equilibrated in buffer E, for 30 minutes at room temperature. After washing to remove excess UAP56 and ATP, beads were resuspended in buffer E and aliquoted. LENG8-PS^M^ or SAC3D1-PS^M^ (2.2 µM / 0.44 µM) was added, followed by a 10-minute incubation at room temperature. Unbound proteins were removed by sequential washes in high-salt buffer (buffer E with 500 mM KCl) and buffer E. RNA-bound proteins were eluted using 0.4 μg benzonase in buffer E for 10 minutes at room temperature, followed by SDS-PAGE analysis and quantification of remaining RNA-clamped UAP56 in Fiji.

### Grating Coupled Interferometry (GCI)

GCI measurements were conducted using a Creoptix WAVE system (Creoptix AG, Switzerland) with 4PCP WAVEchips (quasi-planar polycarboxylate surface). Chips were conditioned in borate buffer (100 mM sodium borate pH 9.0, 1 M NaCl) before immobilization of a monoclonal anti-V5 antibody (R960252, Invitrogen; 2 μg/mL in 10 mM sodium acetate pH 5.0) via amine coupling. The surface was then passivated with 0.5% BSA (in 10 mM sodium acetate pH 5.0) and quenched with 1 M ethanolamine pH 8.0. V5-tagged LENG8-PS^M^ or SAC3D1-PS^M^ complexes were captured to the desired density. UAP56 was injected as a 1:2 dilution series, starting at 5 µM, with or without 200 µM 15U RNA, in 25 mM HEPES pH 7.9, 50 mM KCl, 1 mM MgCl2, 1 mM TCEP, with and without 1 mM ATP at 25°C. Blank injections were used for double referencing, and a DMSO calibration curve corrected for bulk refractive index effects. Data were processed using Creoptix WAVEcontrol software, applying X/Y offset correction, DMSO calibration, and double referencing. A one-to-one binding model was used for fitting, and results were plotted in R.

### ATPase assay

Steady-state ATPase activity of UAP56 was measured using an NADH-coupled enzymatic assay. Final reaction mixtures contained 5 U/mL rabbit muscle pyruvate kinase (Type III, Sigma-Aldrich), 5 U/mL rabbit muscle L-lactic dehydrogenase (Type XI, Sigma-Aldrich), 500 µM phosphoenolpyruvate, and 50 µM NADH. Reactions (10 µL) were assembled in 1536-well plates using buffer F (25 mM HEPES pH 7.9, 40 mM KCl, 0.5 mM MgCl2, 5% glycerol, and 0.5 mM ATP), with either 2 µM UAP56 or 0.1 µM UAP56 in the presence of LENG8–PS^M^ or SAC3D1–PS^M^, and 100 µM 15U RNA when indicated. The decrease in NADH fluorescence emission was monitored at 37°C using a PHERAstar FS plate reader (BMG LABTECH). A calibration curve from a NADH dilution series (0.03–100 µM) was used for quantification. ATPase activity was determined by linear regression of the NADH decay curves, corrected for ATP consumption, and expressed as ATP hydrolysis rates (molecules of ATP hydrolyzed per second per enzyme). Reaction components were analyzed by SDS-PAGE (4–12% gradient) and visualized using Coomassie staining.

### CryoEM sample preparation, imaging, and analysis

*<u>CryoEM sample preparation:</u>* LENG8–PCID2-UAP56-N-UBM–SEM1 (at 0.5 mg ml-1) or SAC3D1– PCID2-UAP56-UCM1-N-UBM–SEM1 (at 0.5 mg ml-1) were incubated in buffer G (25 mM HEPES pH 7.9, 5% glycerol, 1 mM MgCl2, 1 mM TCEP, 100 μM 15U RNA, 1 mM AMP-PNP) on ice for 10 min. CryoEM grids were then prepared by applying 4 µl of the sample to glow-discharged Cu R1.2/1.3 200-mesh holey carbon grids (Quantifoil). Grids were prepared blotted at 8 °C and 90% humidity and plunged into liquid ethane using a Leica EM GP2.

*<u>Cryo-EM data acquisition and processing of a UAP56-LENG8–PS^M^ complex:</u>* Data collection was performed on a Titan Krios G4 electron microscope operating at 300 kV, equipped with a cold field emission gun, a Selectris energy filter (5 eV slit width, ThermoFisher), and a Falcon 4i direct electron detector (ThermoFisher). The objective aperture was retracted, and a 50 µm C2 aperture was used. A total of 5,405 micrographs were recorded using EPU software in .eer format, a pixel size of 0.575 Å/pixel, a total electron dose of 50 e⁻/Å², and defocus values ranging from −1 to −2.5 µm. On-the-fly preprocessing, including motion correction and contrast transfer function (CTF) estimation, was performed using the CryoSPARC Live v113 workflow. Approximately 1.3 million particles were picked in WARP, extracted with a 400 Å box, binned to 1.8 Å/pixel, and subjected to 2D classification. The resulting 2D classes exhibited significant preferred orientation, hindering further 3D reconstruction efforts.

*<u>Cryo-EM data acquisition and processing of a UAP56–SAC3D1–PS^M^ complex:</u>* We collected three datasets with the same microscope specifications and settings as for UAP56-LENG8–PS. Dataset 1 consists of 11,743 micrographs, dataset 2 of 6,006 micrographs collected at a tilt angle of 30° and dataset 3 contains 4,543 micrographs. We again performed on-the-fly preprocessing (patch motion correction and CTF estimation) using the CryoSPARC^113^ live routine before picking 4.5/1.4/0.5 mio particles (dataset 1/2/3) in WARP. For processing in CryoSPARC, particles were extracted with a 400 Å box and binned to 1.8 Å pixel^−1^. After 2D classification we obtained 276,000/47,000/93,000 UAP56– SAC3D1–PS^M^ particles and conducted three rounds of heterogeneous refinement using ab initio models generated with the particles from dataset 1. The resulting 129,495 particles were then re-extracted with a 400 Å box and binned to 0.90 Å pixel^−1^ and subjected to a non-uniform refinement yielding the 3.0 Å UAP56–SAC3D1–PS complex Map A. A further local refinement using a UAP56 mask resulted in the 2.6 Å UAP56–AMP-PNP–RNA Map B.

*<u>Model building for the TREX-2^M^ complex and the UAP56-UCM-1-N-UBM–TREX-2^M^ complex:</u>* Structural modeling of the UAP56–SAC3D1–PS complex began with an Alphafold2 Multimer prediction. The predicted model was docked into the 3.0 Å density (Map A) and manually adjusted using COOT and ISOLDE. Final refinements were performed in Phenix using the phenix.real_space_refine protocol, applying secondary structure and rotamer restraints to optimize fit and stereochemistry.

### HeLa cell culture and cell line generation

HeLa Kyoto cells were grown in Dulbecco’s modified Eagle’s medium (DMEM) supplemented with 10% fetal bovine serum (FBS) and 1% penicillin / streptomycin at 37°C, 5% CO2. Transient transfections were performed using Lipofectamine 3000 (Invitrogen), according to the manufacturer’s instructions. CRISPR/Cas9 mediated genomic knock-ins (KI) using homology dependent repair (HDR) donor vectors^2^ of C-terminal 3xFLAG, 2xHA-FKBP12^F36V^-V(dTAG)^3^ in HeLa Kyoto cells was carried as described before^4^ with guide RNAs (sgRNAs) and homology arms generated using primers listed in Supplementary table 6 and cloned into tagging cassettes carrying Hygromycin or Neomycin resistance genes (plasmids listed in Supplementary table 6). After transfection and antibiotic selection single cell clones were grown and tested by genomic PCR with primers flanking the insertion region, as well as with western blotting analysis. In the GANP-2xHA-dTAG cell line, we observed an additional band, which we interpreted as a truncated protein isoform localized to the cytoplasm. This isoform is produced from an RNA transcript that uses an early polyadenylation site appearing upstream of the tag insertion position. Since dTAG^V^-1 treatment led to rapid and substantial reduction of full length GANP we opted to employ this cell line.

To generate stably expressing LENG8- and ZFC3H1-3xFLAG constructs of WT and mutant variants, HeLa cells were transfected with pBAC vectors as described. Human LENG8 and ZFC3H1 cDNA constructs were cloned and inserted into piggyBAC (pBAC) vectors^5^ using NEBuilder HiFi DNA assembly (NEB). The LENG8 CDS was inserted into a DOX-inducible pBAC vector, harboring a C-terminal 3x-FLAG tag and a Puromycin (PURO) selection marker. The ZFC3H1 CDS was inserted into a constitutively expressed pBAC vector, harboring a C-terminal 3x-FLAG tag and a Blasticidin (BLAST) selection marker. Generated constructs are listed in Supplementary table 6. LENG8- and ZFC3H1-2xHA-dTAG cells were transfected with the pBAC vectors along with a piggyBAC transposase expressing vector (pBASE) with Lipofectamine 3000. Cell pools were selected with PURO or BLAST for 7-10 days until negative control cells died. For induction of expression of LENG8 pBAC constructs, cells were incubated for 24 hours in culture medium supplemented with 1 mg/mL Doxycycline (Sigma-Aldrich) before harvest. Expression of the constructs was validated by western blotting analysis using antibodies against ZFC3H1 or FLAG. Depletion of endogenous dTAG-tagged proteins was performed by the addition of dTAGV-1 to the culture medium for indicated time points at a concentration of 500 nM. Induction of expression of exogenous LENG8-FLAG constructs was performed by adding 1ug/ml doxycycline.

### Immunoprecipitation and analysis of protein complexes

*<u>Western blotting analysis of whole cell extracts</u>*: Whole cell protein lysates were prepared using lysis buffer (20 mM Tris-HCl, 0.5% NP-40, 150 mM NaCl, 1.5 mM MgCl2, 10 mM KCl, 10% Glycerol, 0.5 mM EDTA, pH 7.9) freshly supplemented with protease inhibitors (Roche). Samples were clarified by centrifugation at 20,000 rcf for 10 min. Sample concentrations were adjusted after Bradford measurement and denatured by the addition of NuPage Loading Buffer (Invitrogen) and NuPage Sample Reducing Agent (Invitrogen) before boiling at 95°C for 5 min. SDS-PAGE was carried out on NuPage 4%–12% Bis-Tris (Invitrogen) gels migrated in NuPage MOPS Running Buffer (Thermo) and transferred onto PVDF membranes in NuPage Transfer buffer (Thermo) at 4°C, 15 V overnight. Western blotting analysis was carried out according to standard protocols with the antibodies listed in the Supplementary table 6 and HRP-conjugated secondary antibodies (Agilent). Bands were visualized by Super Signal West Femto chemiluminescent ECL (Thermo) and captured using an ImageQuant 800 imaging systems (GE Healthcare).

*<u>IP followed by western blotting analysis:</u>* Approximately 2×10^7^ cells per IP were extracted in lysis buffer (20 mM Tris-HCl, 0.5% NP-40, 150 mM NaCl, 1.5 mM MgCl_2_, 10 mM KCl, 10% glycerol, 0.5 mM EDTA, pH 7.9) freshly supplemented with protease inhibitors and cleared by centrifugation at 20,000rcf for 20 min. Clarified lysates were incubated overnight at 4°C with anti-FLAG antibody and Protein G Dynabeads (Thermo). Beads were washed 3 times with HT150 extraction buffer, transferring beads to a fresh tube on the final wash. For benzonase treated IPs, samples were resuspended in HT150 buffer freshly supplemented with protease inhibitors and 2 mM MgCl2 and split in two. One half of each sample was mock treated and the other incubated with 500 units of benzonase for 20 min at 25°C / 12000 rpm. Samples were washed twice for 5min at RT in 20mM TrisHCl pH 8 freshly supplemented with 2 mM CaCl2. Proteins were eluted by boiling in 1x NuPage loading buffer (Invitrogen) for 5 min. Supernatants were mixed with 10x Reducing Agent (Invitrogen) and denatured for a further 5 min at 95°C before proceeding with western blotting analysis.

*<u>Immunoprecipitations followed by mass spectrometry:</u>* All IPs were performed label-free and in triplicates. GANP-3xFLAG, PCID2-3xFLAG, SAC3D1-3xFLAG or LENG8-mAID-3xFLAG, and control HeLa Kyoto cells were harvested as described above. Protein extractions were performed using material from 15 million cells per IP with 1 ml extraction buffer (20 mM Tris-HCl, 1% IGEPAL, 150 mM NaCl, 1.5 mM MgCl_2_, 10 mM KCl, 10% Glycerol, 0.5 mM EDTA, pH 7.9) supplemented with 1x protease inhibitors cocktail (Roche). After brief sonication (3 x 10s, Amplitude 1, Branson Sonifier 250), the protein extracts were clarified by centrifugation (20,000rcf for 10min at 4°C). Anti-FLAG magnetic beads were prepared with anti-FLAG M2 antibodies (Sigma F3165) conjugated to Dynabeads M-270 Epoxy (Invitrogen) as previously described^6^. Beads were washed three times with lysis buffer, for endogenous GANP-, LENG8- and PCID2-3xFLAG IPs lysis buffer with additional NaCl to 450 mM final concentration was used. For nuclease treatment beads were resuspended in 40μl extraction buffer with 2 mM MgCl_2_, containing either 1μl Pierce Nuclease (Sigma E1014), benzonase (Sigma) or 1 μl of 1 mg/ml BSA (as indicated for the different experiments) and incubated with agitation at 25°C for 20 min. Beads were washed with extraction buffer once and then proteins were eluted with SDS buffer (2% SDS, 100mM Tris pH 6.8, 10% glycerol) at 25°C, shaking for 5 min. Milder lysis and wash conditions using HT150 buffer (20 mM HEPES pH 7.4, 150 mM NaCl, 0.5% Triton X-100) were applied for 3xFLAG immunoprecipitations of ZFC3H1—both endogenously and exogenously expressed—as well as for exogenously tagged LENG8-3xFLAG. MS sample preparations were performed with the protein aggregation capture (PAC) procedure with proteolytic digestion on MagResyn HILIC beads using trypsin or chymotrypsin as indicated. The peptides were purified and concentrated on C18 stage tips before subjected to LCMS analysis with an Easy nanoLC system coupled directly to a Thermo Scientific Orbitrap Exploris 480 mass spectrometer. MS data were acquired by Data Dependent Acquisition (DDA) and searched against the UniProt protein sequence database using MaxQuant, with “match between runs” and “Label-free quantification” enabled. The MaxQuant protein group output was analyzed with DEP package as previously described^7,8^.

*<u>Chemical fractionation of HeLa cells:</u>* Chemical fractionation was performed using the protocol adapted from^9^. In brief, cells harvested using trypsin digestion were first lysed using cytosol extraction buffer (0.15% NP-40, 10 mM Tris pH 7.4, 150 mM NaCl). Then nuclei were separated from cytoplasmic fractions using centrifugation, followed by nuclei washes using PBS solution and extraction of protein using lysis buffer extraction buffer (20 mM Tris-HCl, 1% IGEPAL, 150 mM NaCl, 1.5 mM MgCl_2_, 10 mM KCl, 10% Glycerol, 0.5 mM EDTA, pH 7.9) or RNA using TRIzol Reagent according to manufacturer’s instructions.

### Immunofluorescence and colocalization analysis

Cells seeded on microscope coverslips were fixed with 4% paraformaldehyde in PBS for 20 min at RT, washed twice with PBS, and permeabilized with 0.1% Triton X-100 in PBS for 10 min at RT. Subsequently, cells were washed with PBS twice and blocked with 5% BSA in PBS-T for 1 hour at RT. Coverslips were incubated for 1 hour at RT with primary antibody dilution in 1% BSA, followed by three 5 min washes with PBS. Then, coverslips were incubated in a secondary antibody dilution with 1% BSA for 1 hour at RT. Finally, cells were washed three times for 5 min with PBS, counterstained with DAPI and mounted onto glass slides using ProLong Gold Antifade Mountant. Images were acquired using a Zeiss LSM 980 confocal microscope equipped with Airyscan 2 under ×40 or ×63 oil-immersion Plan-Apochromat objectives. All images within the same experiment were taken with the same excitation power and exposure time and processed similarly using ZEN Blue 3.6 software. All antibodies and applied concentrations are listed in Supplementary table 6. Pixel-based colocalization analyses were performed using the ZEN 3.6 (blue edition) colocalization module, with threshold setting based on the control background images and extracting the weighted colocalization coefficients for each image. For each cell line, the colocalization coefficient was calculated from six ×40 images in two independent experiments, with at least 139 cells in total included in the analyses.

### RNA extraction and RT-qPCR

HeLa, LENG8-, ZFC3H1- and GANP-2xHA-dTAG cells were treated with 500 nM of dTAG^V^-1 or untreated for 4 hours. RNA was extracted using TRIzol (Invitrogen) and treated with TURBO DNase (Invitrogen) according to the manufacturer’s protocol. For measuring ‘RNA levels’, reverse transcription was carried out with SuperScript III reverse transcriptase (Invitrogen) using 1 µg RNA and a mixture of 20 pmol random hexamer in a 20 µl reaction at 50 °C according to the manufacturer’s protocol. Subsequently, qPCR was performed using Platinum SYBR Green qPCR SuperMix-UDG (Invitrogen) in a ViiA 7 Real-Time PCR machine (Life Technologies with the primers listed in Supplementary table 6). Relative quantities were calculated by normalizing samples to GAPDH mRNA levels. For pA^+^ RNAseq, RNA was quality checked on an Agilent 2100 Bioanalyzer (Agilent Technologies) for integrity before shipping to the sequencing provider.

### pA^+^ RNAseq library generations

All library construction and sequencing were paid services from the Beijing Genome Institute (BGI) in case of total pA^+^ RNAseq and from Lexogen in case of the fractionated and exogenously expressing ZFC3H1 pA^+^ RNAseq. Total RNA was extracted using TRIzol reagent according to manufacturer’s instructions and transferred to BGI or Lexogen, which performed pA^+^ RNA selection using oligo-dT beads followed by strand-specific library preparation and sequencing.

### Analysis of RNAseq data

*<u>Annotation of prematurely terminated transcripts (PTTs)</u>*: Polyadenylated PTTs, displaying sensitivity to ZFC3H1 and/or LENG8 depletion, were annotated using a custom pipeline. Briefly, starting from our transcriptome annotation of HeLa cells^10^, TUs were filtered to be longer or equal to 10 kb. At these TUs, the pA^+^ RNA-seq coverage for the ZFC3H1-2xHA-dTAG and LENG8-2xHA-dTAG cell lines treated with DMSO or dTAG, was measured from TSS to TES with a bin of 50 bp using rtracklayer^11^. Gene bodies were then scaled to 2 kb, replicates averaged, a pseudo-count of 1 added and the log_2_ fold-change (LFC) dTAG/Mock performed for each cell line. Using data from each cell line separately, TUs were then filtered to display increased signal (LFC > 0.2) within the first 200 bp of the scaled gene body and no such difference in the last 200 bp (LFC < 0.2). Of note, the lenient LFC reflected the accumulation of a PTT overlapping with the full TU, and the present criteria filtered out cases where both the full-length TU and the PTT displayed sensitivity to the specific depletions. Following this, the PTT-harboring TUs, identified in ZFC3H1 and LENG8 depletions, were pooled. For each of these the maximum LFC value within the first 200 bp was defined and the last bin reaching 80% of this value along the scaled gene body was used to define an area to screen for the PTT TES. For each TU, in the defined region +/− 5% of the TU length, we measured, without binning or gene body scaling, the coverage from 3’end RNA-seq, non EPAP treated data from ZFC3H1-mAID cells mock or AID treated^4^. LFCs were measured as before and 3’end peaks with ZFC3H1-sensitivity (LFC > 1) over the areas of interest were called. In each area, the strongest peak was considered as the PTT TES. A manual curation of the identified PTTs was then performed to filter out artifacts.

*<u>Processing of total and fractionated pA+ RNA seq data:</u>* The raw sequence reads received from service providers were first quality checked using FastQC (Babraham Bioinformatics group). Reads were then trimmed for adaptors and filtered using Trim Galore (Babraham Bioinformatics group). Trimmed reads were mapped to hg38, using Hisat2 in paired-end mode^14^. Mapped files were sorted and checked for pairing using SAMtools^15^. Reads were then deduplicated using MarkDuplicates^16^ (Picard – Broad Institute) and further filtered to keep only unique mappers by using SAMtools. Relative samples size was then estimated by generating coverage counts using htseq-count^17^ (HTSeq-counts) over the Gencode annotation to avoid any bias due to accumulation of short unstable transcripts present in our in-house annotation, and then analyzed by DESeq2 (ref.^18^) to define size factors. In the case of fractionated pA^+^ RNAseq, size factors were measured separately for the nuclear and cytoplasmic fractions to avoid any compensation of compartment specific phenotypes. Finally, reads were converted to bigwig (bw) files normalized to size factors using bamCoverage (Deeptools)^19^. The RNA sample of HeLa 4-hour dTAG^V^-1 replicate 4 from the nuclear fraction appeared to suffer from a strong technical issue arising from large ribosomal RNA contamination. This replicate was therefore eliminated from all analyses but still listed as part of GEO dataset.

*<u>Differential expression analysis:</u>* RNA sensitivities to LENG8-, ZFC3H1- and GANP-depletions were all defined based on DESeq2 differential expression analysis of total pA^+^ RNAseq using untreated cells as controls. For each depletion TUs with adjusted p-values < 0.1 in DESeq2 (ref.^18^) analysis were considered as measurable and log_2_ fold change (LFC) over control > 0.5 was counted as up-regulated, while the LFC coverage over control < −0.5 was counted as downregulated. Due to the strong correlation between coverage changes of LENG8- and ZFC3H1-depletions (Fig. 3c), ‘PAXT sensitive’ TUs were defined as upregulated in either of the two depletions. Plots exploring the relationship between exons or processed RNA lengths and PAXT sensitivites (Fig. 3e, Supplementary Fig. 5k, Supplementary Fig. 7i-j) were based on our published in-house HeLa transcript annotation^10^ and LFC coverage for all TUs with adjusted p-values < 0.1 in DESeq2.

*<u>Nuclear to cytoplasmic ratios measurements:</u>* Nuclear to cytoplasmic ratios were calculated for each TU using non−log transformed counts of nuclear and cytoplasmic pA^+^ RNA coverages. Zero value counts were filled with minimal values.

*<u>TU clustering based on fractionated pA^+^ RNAseq behavior:</u>* First the average LFC coverage, as measured by rtracklayers, was calculated for the total, nuclear and cytoplasmic fractions. For the fractionated sequencing, all proteins depletions were compared to the maternal HeLa cell line treated with dTAG^v^-1. The LFC upon ZFC3H1 depletion was then used separately in the nuclear and cytoplasmic fractions, at each TU, to define a behavior as ‘up’ (> 0.5), ‘down’ (< −0.5), or ‘unaffected’. Nine clusters were then generated corresponding to all possible combinations (nuclear ‘up’/cytoplasmic ‘up’, nuclear ‘up’/cytoplasmic ‘unaffected’ etc…). Small clusters (with less than 200 TUs) were removed from the final heatmap in Supplementary Fig. 6c.

*<u>Analysis of transcripts with retained introns:</u>* For every intron unspliced reads spanning the 5’ and 3’ splice sites were counted using custom code relying on Samtools^15^, and every intron with at least one unspliced read at both junctions in unperturbed HeLa cells was considered as retained. The genomic coordinates of detained introns were obtained from (Boutz et al.^19^). As this annotation originates from four combined cell lines, we first merged it with our in-house HeLa specific annotation^10^. Considering the generally unspliced nature of detained introns, we first filtered out these when overlapping totally or partially introns from our annotations. We then further filtered detained introns to be fully included in our exons to avoid overhang at the TSS or TES due to alternative isoforms. Finally, the few cases where a detained intron was starting or ending a TU, without being preceded or followed by an exon, were filtered out. Similarly, reads spanning 5’ and 3’ splice junctions for both retained and detained introns were counted in total, nuclear and cytoplasmic fractions. Introns were dual splice junctions showed an increase upon ZFC3H1- or LENG8-depletion were counted as PAXT-sensitive.

*<u>Metagene profiles, heatmaps and display of sequencing information:</u>* Metagene profiles and heatmaps were produced using custom R and Python scripts. Briefly, the rtracklayer R package was used to collect read coverage values for the window −500 bp / +500 bp according to the TSS or TES, or over specific exonic/intronic features. Coverage values were then binned in 50 nt bins and log_2_-transformed after the addition of a pseudo-count of 1. This measurement of coverage was then used to compute LFC values and generate subsequent plots. Heatmaps were made using custom R or Python code based on the R package ComplexHeatmap^21^ or Seaborn^22^, respectively. The mean of coverage values across TUs over each bin were also computed and plotted as metagene profiles using custom R code. A 95% confidence interval of the mean coverage was displayed for each sample and was measured through 50 steps of bootstrap samplings with replacement. Aggregate plots and heatmaps of sequencing data were generated based on BigWig files using customized R scripts. Genome browser views based on BigWig files were generated using the R package seqNdisplayR^23^.

### Statistical data analysis

In addition to the built-in statistical tests provided by software packages such as Zen Blue, DESeq2, and DEP, further statistical analyses were performed using two-sided t-tests or Welch’s t-test when group sizes differed substantially. Pearson correlation coefficients were used to assess the correlation between LFCs following ZFC3H1-, LENG8- or GANP-depletion. Box plots show the mean, interquartile range, and whiskers represent data distribution; statistical significance is indicated directly on the plots. Outlier dots were excluded from the visual display for clarity but were included in the statistical analysis.

### in vivo RNA-binding assays

Cells expressing LENG8-3xFLAG of WT or R563A mutant variants were induced with doxycycline for 24 hours and crosslinked with 150 mJ/cm^2^ of 254 nm UV lamp using Stratalinker2400 (Stratagene). Lysate preparations, anti-FLAG IPs, RNAse I (Thermo Scientific), TurboDNAse treatments (Thermo Scientific), radiolabeling using γ-^32^P ATP (PerkinElmer) and PAGE of RNA-protein complexes were performed as described^24^. The phosphor imaging of gels with radiolabeled samples was performed using a Typhoon scanner (Amersham).

### Puromycin labeling assays

HeLa, LENG8-, ZFC3H1- and GANP-2xHA-dTAG cells were grown in the presence of 500 uM dTAG^V^-1 for additional 0, 4 and 24 hours. Before harvesting by snap freezing, 5 ug/ml of Puromucin was added to cell media. Puromycin incorporation was assessed by western blotting analysis.

### Whole cell proteome analysis using pulsed SILAC

HeLa, LENG8-, ZFC3H1- and GANP-2xHA-dTAG cell lines were initially cultured in DMEM medium in the presence of 73 mg/L l-lysine HCl and 28 mg/L l-arginine HCl, (Sigma) (Lys0/Arg0 medium) for 24 hours. Cells were then pre-treated with either 500 µM dTAG^V^-1 or an equivalent volume of DMSO for 4 hours. Following this, the medium was switched to medium containing 73 mg/L l-lysine HCl and 28 mg/L l-arginine HCl l-lysine (^13^C_6_^15^N_2_) and l-arginine (^13^C_6_^15^N_4_), (for the Lys8/Arg10 medium) with either dTAG^V^-1 or DMSO, and cells were cultured for an additional 24 hours under the same conditions. In parallel, a matched set of cells was maintained in Lys0/Arg0 medium under the same condition (dTAG^V^-1 or DMSO). After treatment, cells were harvested by snap freezing and SILAC sample preparation and MS were carried out as described^25^. SILAC ratios of Lys8/Arg10 *vs.* Lys0/Arg0 peptides were calculated for each sample. To calculate differential protein expression, the DEP package^26^ was used to analyze mean LFQ intensities differences of Lys8/Arg10-labelled peptides.

## Code availability

The article does not report any original code.

## Data availability

Three-dimensional cryo-EM density maps of UAP56–RNA–SAC3D1–PS^M^ have been deposited to the Electron Microscopy Data Bank under the accession numbers EMD-54282 (Composite Map), EMD-54283 (Map-A) and EMD-54284 (Map-B). The coordinate file of UAP56–RNA–SAC3D1–PS^M^ has been deposited to the Protein Data Bank under the accession number 9RV1.

All newly generated RNA-seq data are available at GEO accession code: GSE301785 (reviewer token: ejupugyyxbwtbov)

All newly generated proteomics data are avaliable at PRIDE accession code: PXDxxxx

Raw image files are deposited on Mendeley Data and are available at: https://data.mendeley.com/preview/s73vw4pyh7?a=9b8568ca-f4c2-47ac-b05c-ea865feed433

Plasmid DNA constructs and stable human cell lines, are available from the authors upon request.

## SUPPLEMENTARY FIGURES

**Supplementary Figure 1.**
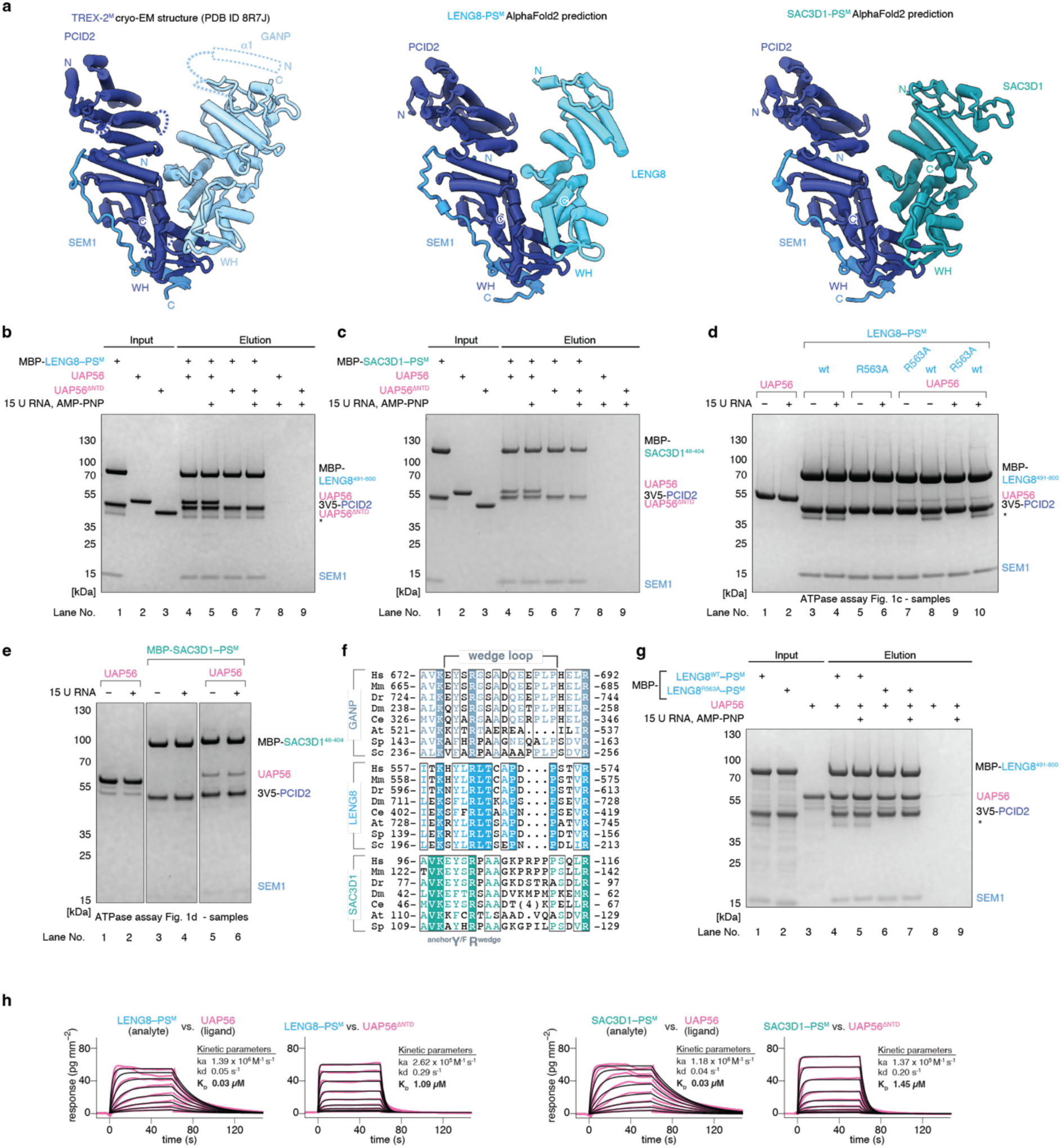
Biochemical characterization of TREX-2-like complexes. **a.** Cryo-EM structure of TREX-2^M^ (PDB ID 8R7J) (ref.^7^) (left) and AlphaFold2 (ref.^21,22^) predictions of LENG8^491–800^–PCID2–SEM1 (center) and SAC3D1^48–404^–PCID2–SEM1 (right) in cartoon representations. PCID2, dark blue; SEM1, blue; GANP, light blue; LENG8, mid blue; SAC3D1, green blue. **b-c.** Coomassie-stained SDS-PAGE gels of *in vitro* pulldown experiments demonstrating that UAP56 interacts with LENG8–PS^M^ (b) and SAC3D1–PS^M^ (c) *in vitro* and that the UAP56 NTD is required for this interaction. Recombinant MBP-LENG8–PS^M^ or MBP-SAC3D1–PS^M^ were immobilized on amylose beads through the MBP tag and incubated with UAP56 or UAP56^ι1NTD^ and with or without 15 nucleotide poly-uridine RNA and AMP-PNP as indicated. After washes, bead-bound proteins were eluted with maltose containing buffer. *Denotes contaminant. **d-e.**Coomassie-stained SDS-PAGE gels of samples containing the indicated components as used in UAP56 ATPase assays in Fig. 1c (d) and 1d (e). For each reaction condition an aliquot was analyzed by SDS-PAGE and Coomassie-staining. *Denotes contaminant. **f.** Multiple sequence alignment of wedge loop sequences of GANP (top), LENG8 (middle) and SAC3D1 (bottom) proteins from selected model species. Amino acidsI are colored by conservation (colored text, conserved residue; colored background, invariant residue). GANP proteins are from *H. sapiens* (Hs, Uniprot ID O60318), *M. musculus* (Mm, Q9WUU9), *D. rerio* (Dr, F1Q712), D*. melanogaster* (Dm, Q9U3V9), *C. elegans* (Ce, Q19643), *A. thaliana* (At, F4JAU2), *S. pombe* (Sp, O74889) and *S. cerevisiae* (Sc, P46674); LENG8 proteins are from *H. sapiens* (Hs, Uniprot ID Q96PV6), *M. musculus* (Mm, Q8CBY3), *D. rerio* (Dr, A4QNR8), *D. melanogaster* (Dm, B3DNL8), *C. elegans* (Ce, Q8I7L2), *A. thaliana* (At, A0A178VUB3), *S. pombe* (Sp, Q1MTP1) and *S. cerevisiae* (Sc, Q12049); SAC3D1 proteins are from *H. sapiens* (Hs, A6NKF1), *M. musculus* (Mm, A6H687), *D. rerio* (Dr, A0AB13A8D7), *D. melanogaster* (Dm, Q9VSZ4), *C. elegans* (Ce, Q8WQE7), *A. thaliana* (At, Q67XV2) and *S. pombe* (Sp, Q9USI4). **g.** *In vitro* pulldown assay as in b, but using the LENG8^R563A^ wedge loop mutant. **h.** Grating-coupled interferometry (GCI) assays employing LENG8–PS^M^ (top) or SAC3D1–PS^M^ (bottom) complexes immobilized as the analyte on a microfluidic chip and with UAP56 (left), or its UAP56^ι1NTD^ variant (right), flown in at increasing concentrations as the ligand. Shown are sensograms (pink lines), the fitted model (black lines) and a binding kinetics summary table.

**Supplementary Figure 2.**
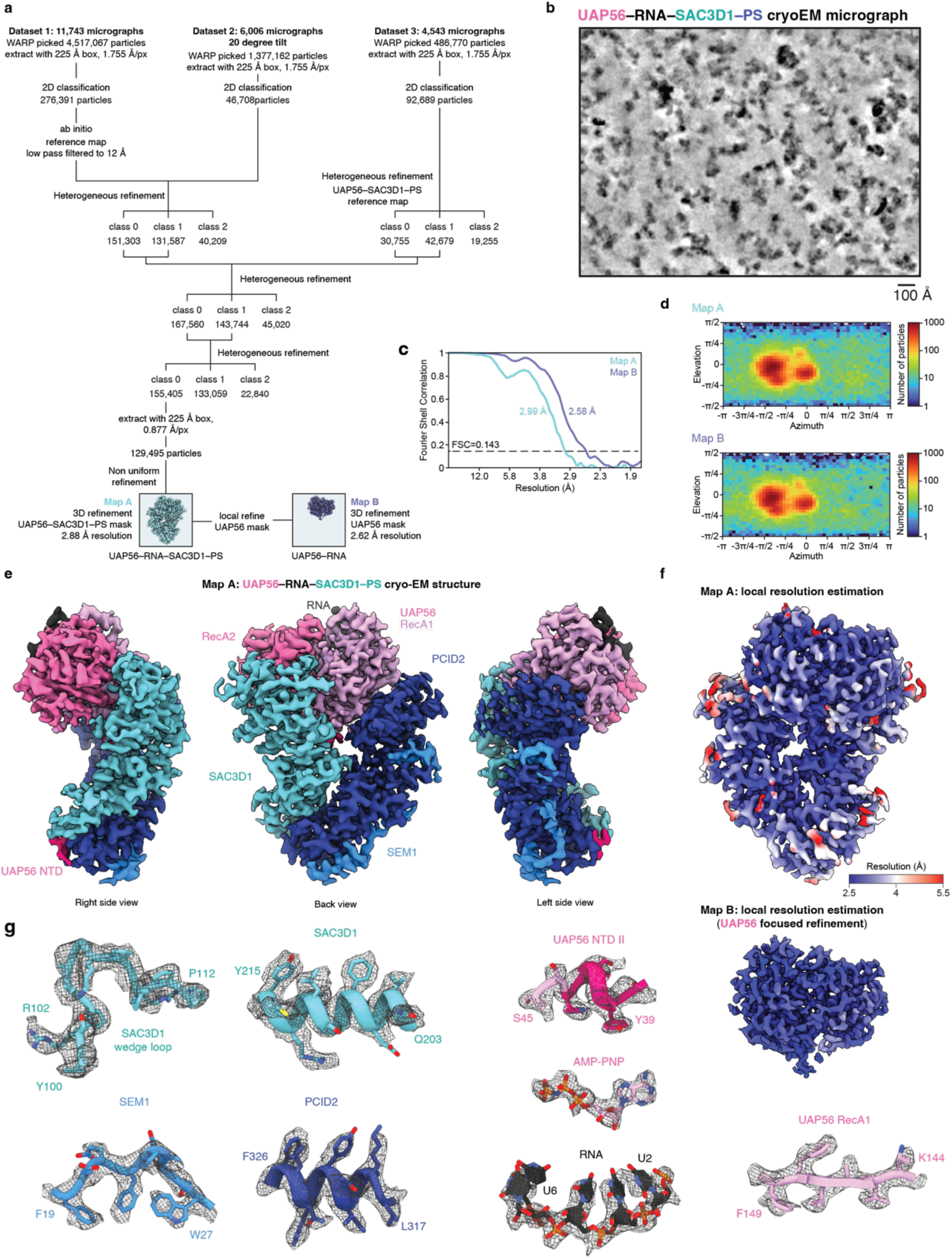
UAP56–RNA–SAC3D1–PS cryo-EM analysis. **a.** Three-dimensional image classification tree. From three datasets, containing 11,743, 6,006 (collected at 20-degree stage tilt) and 4543 micrographs, 4,517,067, 1,377,162 and 486,770 particles were picked through a BoxNet in WARP and processed in cryoSPARC. The final particle stack contained 126,495 particles and was refined to 3.0 Å resolution for the UAP56–RNA–SAC3D1–PS complex (Map A). A focused refinement using a mask comprising UAP56 and the bound RNA yielded a 2.6 Å density of UAP56–AMP-PNP–RNA (Map B). **b.** Denoised cryo-EM micrograph of the UAP56–RNA–SAC3D1–PS sample. Scale bar, 100 Å. **c.** Gold-standard Fourier shell correlation (FSC = 0.143) of the UAP56–RNA–SAC3D1–PS complex (Map A), and UAP56–AMP-PNP–RNA (Map B) cryo-EM maps. **d.** cryoSPARC orientation distribution plots for UAP56–RNA–SAC3D1–PS (Map A), and UAP56– AMP-PNP–RNA (Map B) cryo-EM maps. **e.** UAP56–RNA–SAC3D1–PCID2–SEM1 cryo-EM density (Map A) as in Fig. 1g, colored by subunit identity. SEM1, blue; PCID2, dark blue; SAC3D148-404, green blue; UAP56, shades of pink; RNA, black. **f.** UAP56–RNA–SAC3D1–PS (Map A), and UAP56–AMP-PNP–RNA (Map B) cryo-EM maps colored by local resolution. **g.** Representative segments from Map A (SAC3D1, PCID2, SEM1, UAP56 NTD) and Map B (UAP56, RNA, AMP-PNP) in cartoon representation with side chains shown as sticks and superimposed on the respective cryo-EM densities.

**Supplementary Figure 3.**
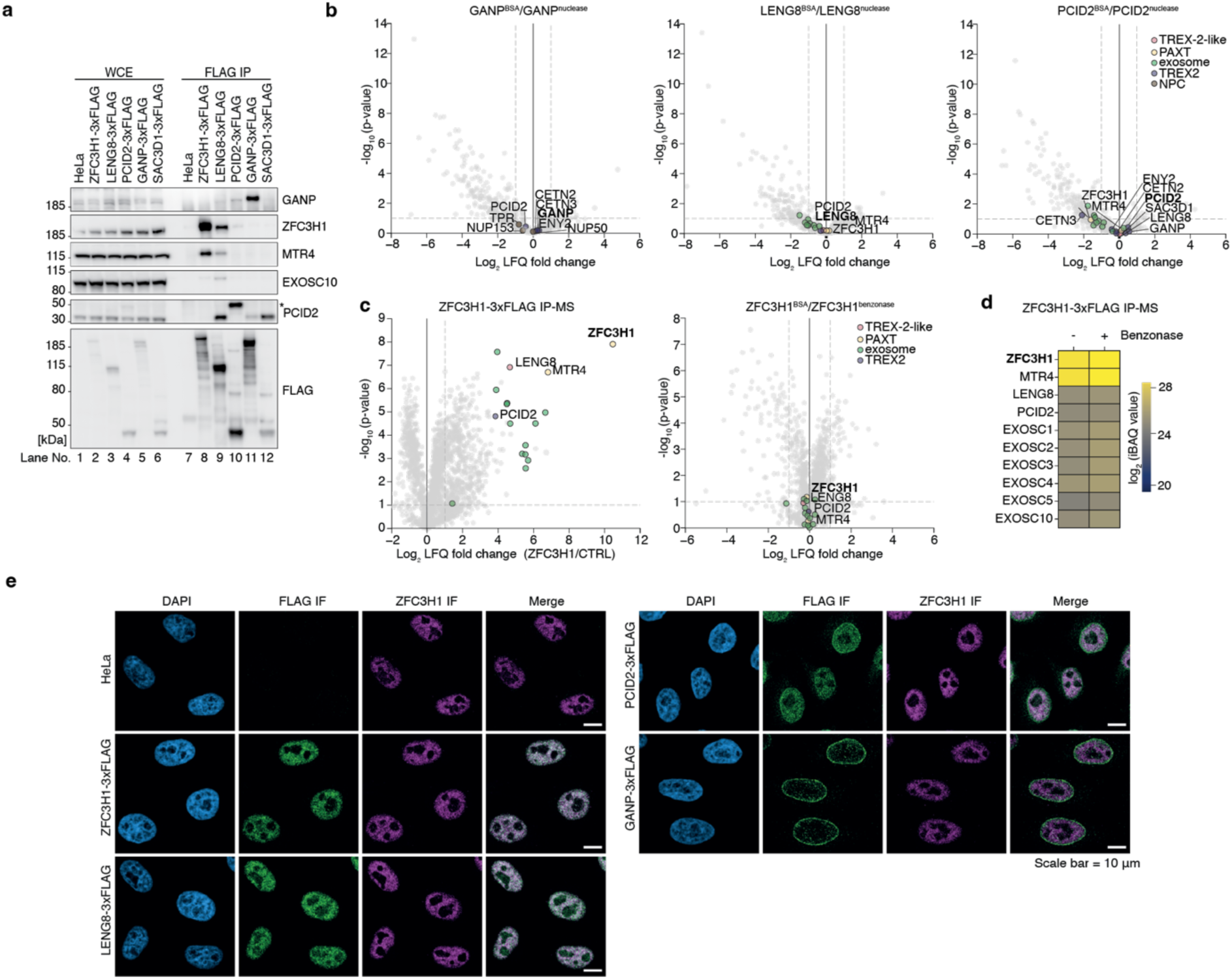
Multiple TREX-2-like complexes in human cells. **a.** Western blotting analysis of whole cell extracts (WCE, left part) or eluates after anti-FLAG IP (right part) of extracts from endogenously 3xFLAG-tagged ZFC3H1, LENG8, PCID2, GANP or SAC3D1 cells. Maternal HeLa cells were included as a negative control. Asterisk denotes the 3xFLAG-tagged PCID2 band on the anti-PCID2 blot. Membranes were probed with GANP-, ZFC3H1-, MTR4-, EXOSC10-, PCID2- and FLAG-specific antibodies as indicated. **b.** Volcano plots related to Fig. 2b, but showing log_2_ fold changes of LFQ values of interactors in Pierce universal nuclease-treated relative to non-treated IP samples of the indicated bait proteins (x-axes). **c.** Volcano plot as in Fig. 2b (left panel) or Supplementary Fig. 3b (right panel), but of IP-MS analysis of extract from ZFC3H1-3xFLAG cells. **d.** Heatmap as in Fig. 2c, but of IP-MS analysis of extract from ZFC3H1-3xFLAG HeLa cells, showing Benzonase-resistant enrichments of MTR4, LENG8, PCID2 and exosome subunits. **e.** Immuno-localization analysis as in Fig. 2a. Shown are DAPI (left column)-, anti-FLAG (left-mid column)-and anti-ZFC3H1 (right-mid column)-stained HeLa cell lines, expressing the indicated C-terminally 3xFLAG tagged proteins. Right column shows merges of FLAG and ZFC3H1 signals. Maternal HeLa cells are shown as a negative control for the FLAG IF. Related to Figure 2d.

**Supplementary Figure 4.**
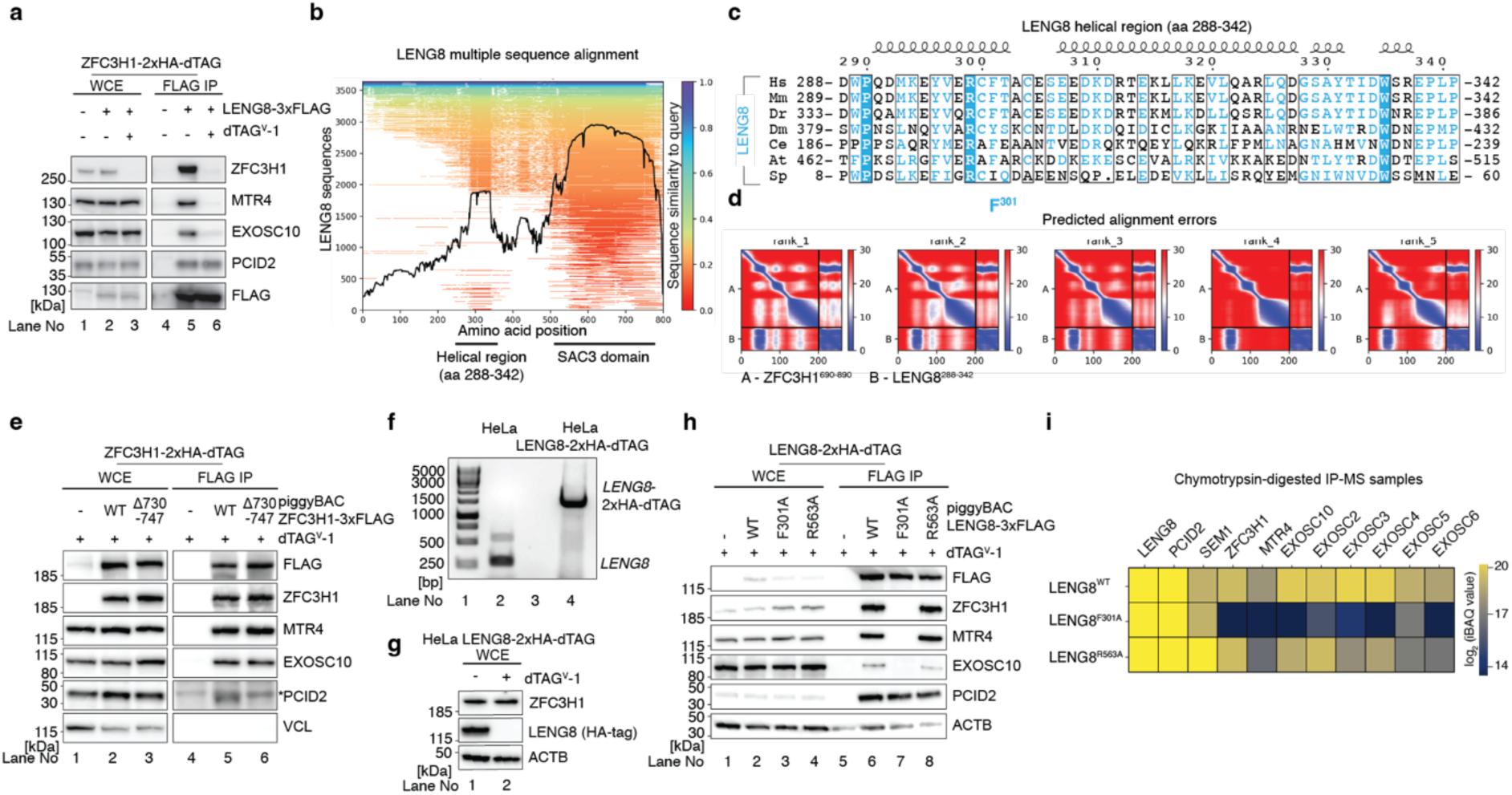
LENG8-PS interaction with PAXT and the exosome is mediated by direct binding of ZFC3H1 and LENG8. **a.** Western blotting analysis of WCE (left part) or eluates of anti-FLAG IP (right part) of extracts from HeLa cells, expressing endogenously dTAG-tagged ZFC3H1, and which were not (-) or transiently transfected (+) with plasmid expressing 3xFLAG-tagged LENG8, without or in presence of dTAGV-1 as indicated. Membranes were probed with ZFC3H1, MTR4, EXOSC10, PCID2 and FLAG-tag-specific antibodies as indicated. **b.** Multiple sequence alignment of *H. sapiens* LENG8 protein with LENG8 sequences from the AlphaFold2 MMseq2 database. Conserved helical regions and SAC3 domains are indicated by black lines. **c.** Multiple sequence alignment as in Supplementary Fig. 1g, but of the conserved region of human LENG8^288–342^. The phenylalanine (F301) critical for interaction with ZFC3H1 is highlighted. **d.** Predicted Alignment Error plots for AlphaFold2 predictions of ZFC3H1^690–890^ and LENG8^288–342^: Alignment errors for the top 5 models are shown. Related to Fig. 2e. **e.** Western blotting analysis as in (**a**), but of extracts from HeLa cells, expressing endogenously dTAG-tagged ZFC3H1 without or with piggyBac-integrated exogenous ZFC3H1^WT^-3xFLAG or ZFC3H1^Δ730–747^–3xFLAG variants as indicated. Membranes were probed with FLAG-tag, ZFC3H1, MTR4, EXOSC10, PCID2 and vinculin (VCL)-specific antibodies as indicated. Asterisk indicates unspecific band. **f.** PCR analysis with amplicon spanning the region of CRISPR-mediated modification of the *LENG8* gene, using DNA from maternal HeLa- or LENG8-2xHA-dTAG cells (see also Methods). **g.** Western blotting analysis of whole cell extracts of LENG8-2xHA-dTAG cells untreated or treated with dTAG^V^-1 for 4 hours. Membranes were probed with HA-tag for LENG8, ZFC3H1, actin-specific antibodies as indicated. **h.** Western blotting analysis of WCE (left part) or of eluates from anti-FLAG IP (right part) from extracts of HeLa cells with endogenously dTAG-tagged LENG8 without or with piggyBac-integrated exogenous LENG8^WT^-3xFLAG, LENG8^F301A^- or LENG8^R563A^-3xFLAG constructs. Membranes were probed with FLAG-tag, ZFC3H1, MTR4, EXOSC10, PCID2 and actin-specific antibodies as indicated. **i.** Heatmap as in Figure 2c but for LENG8^WT^-LENG8^F301A^-, or LENG8^R563A^-3xFLAG IPs from (g). MS was performed using chymotrypsin treated eluates to enable detection of SEM1. iBAQ values from control FLAG IPs in HeLa were subtracted.

**Supplementary Figure 5.**
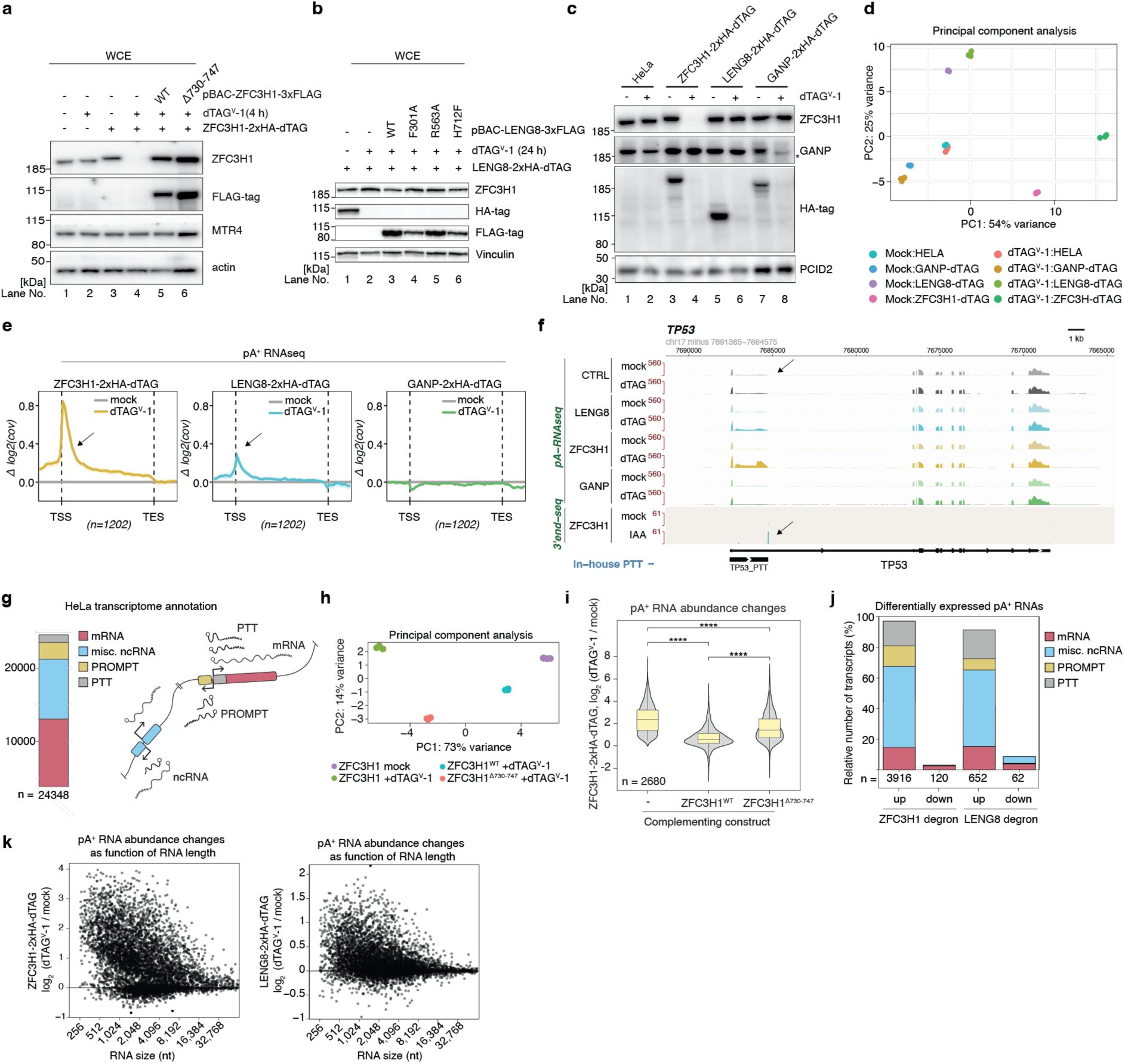
Interaction between ZFC3H1 and LENG8 is required to suppress pA^+^ RNA. **a.** Western blotting analysis of WCEs from maternal HeLa- or ZFC3H1-2xHA-dTAG-cells treated (+), or not (-), for 4 hours with dTAG^V^-1. PiggyBac expression of ZFC3H1^WT^-3xFLAG or ZFC3H1^Δ730–747^–3xFLAG was conducted as indicated. Membranes were probed with ZFC3H1-, FLAG-, MTR4- and actin-specific Abs as indicated. Related to Fig. 3a. **b.** As in (a) but employing LENG8-2xHA-dTAG cells subjected to 24 hours of dTAG^V^-1 treatment (+), or not (-). LENG8^WT^-, LENG8^F301A^-, LENG8R^563A^- or LENG8^H712F^-3xFLAG expression was conducted as indicated. Membranes were probed with ZFC3H1-, HA-, FLAG- and vinculin-specific Abs as indicated. Related to Figure 3b. **c.** Western blotting analysis of WCEs from maternal HeLa- or ZFC3H1-, LENG8- or GANP- 2xHA-dTAG cell lines mock treated (-) or treated for 4 hours with dTAG^V^-1 (+) as indicated. Membranes were probed with ZFC3H1-, GANP-, HA- and PCID2-specific Abs as indicated. Truncated GANP isoform is indicated by an asterisk. **d.** Principal component analysis of obtained biological replicates of indicated pA^+^ RNAseq samples. **e.** Aggregate plots of Δlog_2_(coverage) signals of pA^+^ RNA-seq data from ZFC3H1-, LENG8- and GANP- 2xHA-dTAG cell lines treated for 4 hours with dTAG^V^-1 *vs.* their mock-treated controls. Signals were plotted over TUs expressing identified PTTs and displayed from 500 bp TSS-upstream to 500 bp TES-downstream regions (note arrows pointing to the TSS-proximal area with increased pA^+^ RNA coverage upon dTAG^V^-1 treatment). The TU body (TSS to TES) was normalized to 2 kb for each TU. Data are displayed with 90% confidence intervals (lighter shades around the curves). Numbers (n) of aggregated TUs are indicated. **f.** Genome browser view displaying pA^+^ RNAseq and 3’-RNAseq data, used in the PTT annotation pipeline, over the *TP53* gene with annotated *TP53* and *TP53 PTT* TUs. Arrows point to the TSS-proximal area with increased pA^+^ RNA coverage and to the pA^+^ 3’end of the 3’end-seq data. **g.** Left: Simplified biotype overview of the 24348 annotated RNAs of the HeLa transcriptome. Right: Cartoon representation of the annotated biotypes. **h.** Principal component analysis of obtained biological replicates of indicated pA^+^ RNAseq samples. **i.** Violin plots of pA^+^ RNA log_2_ fold abundance changes of RNAs upregulated upon ZFC3H1 depletion (per DEseq2, see Methods) after dTAG^V^-1 treatment for 4 hours of ZFC3H1-2xHA-dTAG cells not expressing (left), expressing ZFC3H1^WT^ (mid) or expressing the LENG8-binding deficient mutant ZFC3H1^Δ730–747^ (right). Student’s t-test were calculated between conditions (****p-value<0.0001). **j.** DESeq2 analysis of pA^+^ RNAseq data derived from ZFC3H1- and LENG8-2xHA-dTAG cells treated, or not, with dTAG^V^-1 for 4 hours. Biotypes indicated as in panel g. **k.** Scatterplots of pA^+^ RNA log_2_ fold abundance changes after dTAG^V^-1 treatment for 4 hours of ZFC3H1-(left) or LENG8 (right)-2xHA-dTAG cells as functions of log_2_-scaled lengths in the range 256 to 65,536 nucleotides (see also Methods) of the respective processed RNAs. All transcripts with measurable fold changes (−log_10_ (adj. p-value) > 1) from DESeq2 analysis were included.

**Supplementary Figure 6.**
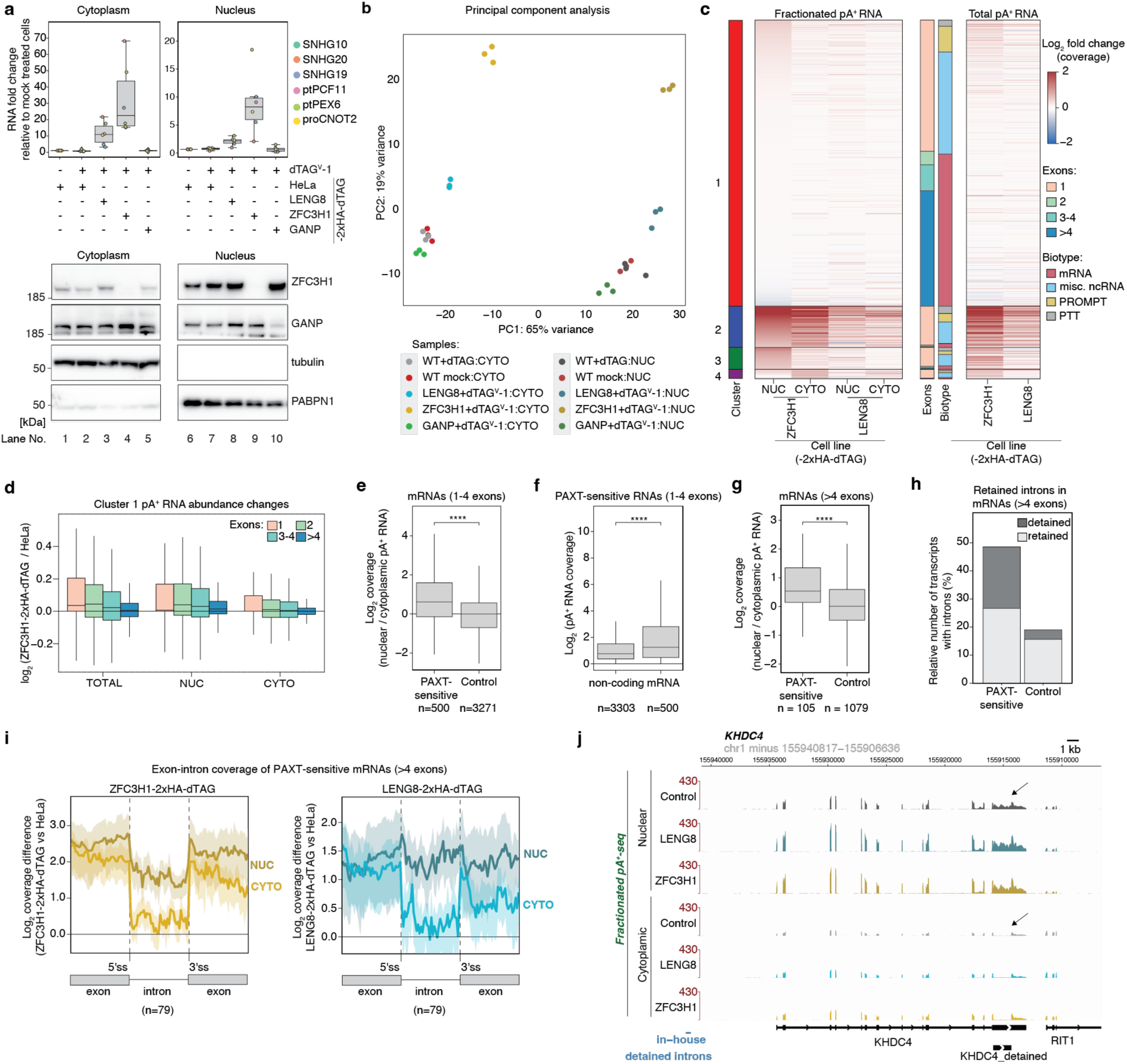
The LENG8-PAXT axis retains nuclear pA^+^ RNAs. **a.** Top: Boxplots of RT-qPCR analysis of select PAXT substrates in nuclear and cytoplasmic fractions of biochemically fractionated maternal HeLa cells or the indicated 2xHA-dTAG-tagged cell lines with dTAG^V^-1-induced depletion of LENG8, ZFC3H1 or GANP for 4 hours. RNA fold changes were calculated relative to values of non-treated HeLa control cells and normalized to GAPDH mRNA, separately for cytoplasmic (left) and nuclear (right) fractions. Dots indicate mean values of three biological replicates for each RNA. Bottom: Western blotting analysis of employed cytoplasmic (left) and nuclear (right) extracts Membranes were probed with ZFC3H1-, GANP-, tubulin- and PABPN1-specific Abs as indicated. **b.** Principal component analysis of obtained biological replicates of indicated pA^+^ RNAseq samples. **c.** Heatmaps of pA^+^ RNA log_2_ fold abundance changes in nuclear (NUC) and cytoplasmic (CYTO) fractions (left panel) or total (right panel) samples of dTAG^V^-1-treated (4 hours) ZFC3H1- or LENG8-2xHA-dTAG cells relative to maternal HeLa cells. Transcripts were clustered based on NUC and CYTO abundance changes upon ZFC3H1 depletion with a cutoff log_2_ fold change of 0.5. Stacked bar plots in between heatmaps display transcript biotypes and contained exon numbers within each cluster. **d.** Boxplots of pA^+^ RNA log_2_ fold abundance changes upon ZFC3H1 depletion for all transcripts in Cluster 1 of panel c. Transcripts were grouped in compartments and color-coded by exon numbers. **e.** Boxplots of log_2_-scaled nuclear-cytoplasmic ratios of pA^+^ RNA log_2_ coverages in unperturbed HeLa cells for low-exon (1-4) PAXT-sensitive mRNAs and control mRNAs of the same exon numbers. Welch’s t-test were calculated between conditions, combining the results for all RNAs tested (**** p-value <0.0001). **f.** Boxplots of pA^+^ RNA log_2_ levels of PAXT-sensitive ncRNAs or mRNAs (1-4 exons) in unperturbed HeLa cells. PAXT-sensitive transcripts were defined as in Fig. 3d. Welch’s t-test were calculated between conditions, combining the results for all RNAs tested (**** p-value <0.0001). **g.** Boxplots of nuclear-cytoplasmic ratios as in panel e, but for PAXT-sensitive mRNAs (>4 exons) compared to control mRNAs of the same expression levels, lengths and contained exon numbers (Methods). Welch’s t-test were calculated between conditions, combining the results for all RNAs tested (**** p-value <0.0001). **h.** Proportion of mRNAs (>4 exons) harboring detained or retained introns in PAXT-sensitive mRNAs compared to control mRNAs of the same expression levels, lengths and contained exon numbers (Methods). **i.** Aggregate plots displaying log_2_-scaled differences (note: not log_2_ fold changes, as exons and introns differ in their coverage ranges) of RNAseq coverage over retained introns, and their spanning exons, of PAXT-sensitive mRNAs (>4 exons) in NUC or CYTO fractions of depleted ZFC3H1 (left) or LENG8 (right) cells as compared to unperturbed HeLa cells. **j.** Genome browser view as in Supplementary Fig. 5f, but showing nuclear and cytoplasmic pA^+^ RNAseq data, over the *KHDC4* gene with its annotated detained intron (depicted by an arrow).

**Supplementary Figure 7.**
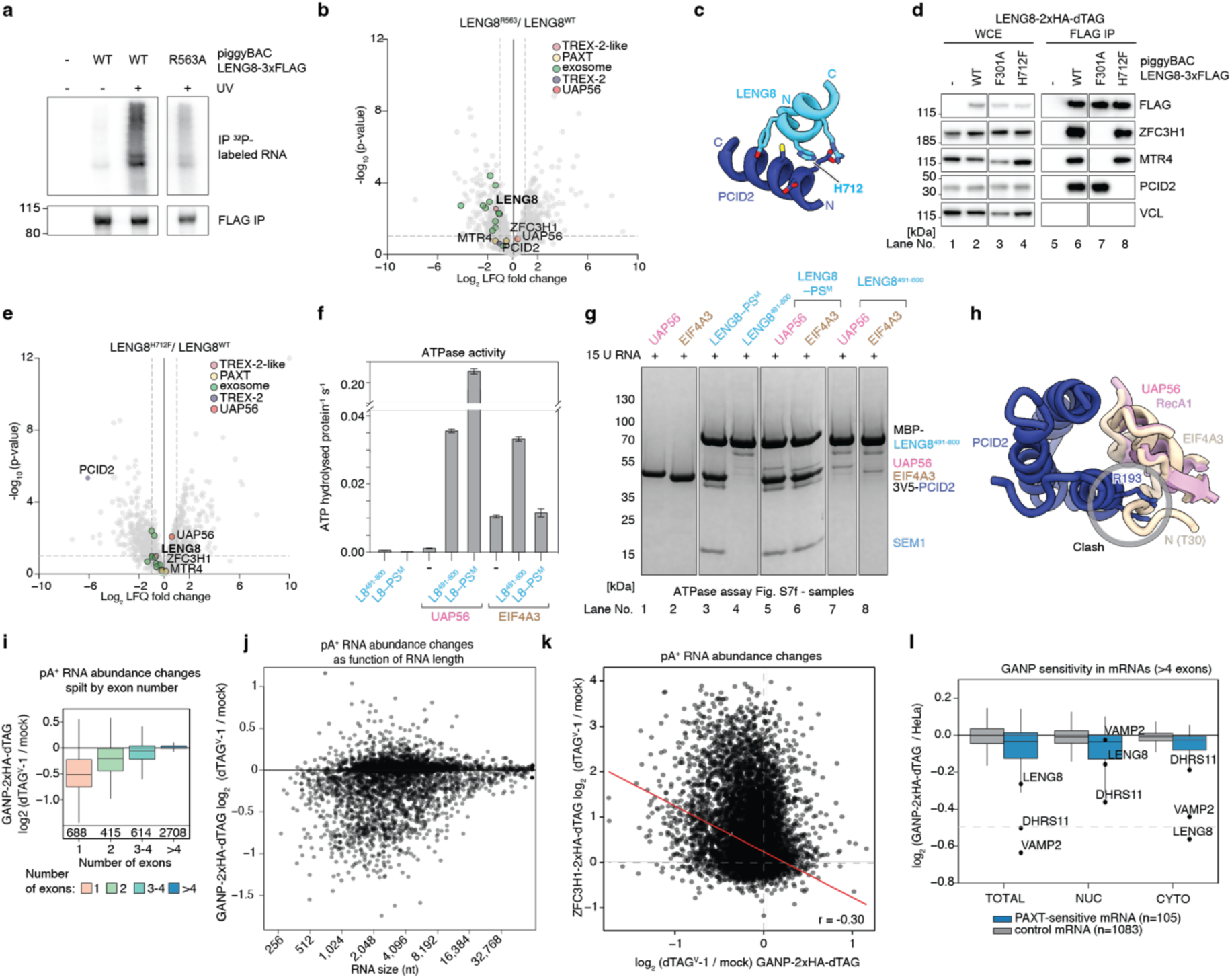
TREX-2 depletion results in downregulation of export-competent PAXT substrates. **a.** Autoradiogram of RNA signals deriving from IPs of LENG8WT or LENG8^R563A^ constructs upon 150 mJ/cm^2^ UV crosslinking (+) or not (-). Western blotting analysis of the FLAG IPs were probed with FLAG Ab (‘FLAG IP’). **b.** Volcano plot as in Fig. 2f, but for LENG8^R563A^-3xFLAG relative to LENG8WT-3xFLAG sample data. UAP56, LENG8, PCID2 as well as PAXT- and exosome-components are color coded. **c.** AlphaFold2 model of interacting regions of LENG8 (blue) and PCID2 (dark blue). The conserved H712 residue of LENG8 is highlighted. **d.** Western blotting analysis of WCEs (left) or anti-FLAG IP (right part) samples of HeLa cells with endogenously dTAG-tagged LENG8 without or with piggyBac-integrated exogenous LENG8WT-3xFLAG, LENG8^F301A^- or LENG8^H712F^-3xFLAG variants. Membranes were probed with FLAG-tag, ZFC3H1, MTR4, PCID2, vinculin (VCL)-specific Abs as indicated. **e.** Volcano plot as in (b), but for LENG8^H712F^-3xFLAG relative to LENG8WT-3xFLAG sample data. **f.** ATPase assay as in Fig. 1b. Plotted are the ATPase rates, in the presence of 15 poly-uridine RNA, of LENG8 or the LENG8–PS^M^ complex in isolation or with either UAP56 or EIF4A3 added. **g.** ATPase assay protein samples as in panel (Supplementary Fig. 1e-f), but for the experiment shown in panel (f). **h.** Cartoon showing PCID2 (dark blue) and UAP56 (pink) as positioned in the SAC3D1-PS^M^ and EIF4A3 from (PDB ID 7ZNJ, chain A, brown). EIF4A3 was superimposed onto UAP56 RecA1 (RMSD = 0.87 Å across 189 atom pairs), revealing clashes between EIF4A3 residues 30-35 and PCID2. **i.** Boxplots of pA^+^ RNA log_2_ fold abundance changes as in Fig. 3e, but for GANP-2xHA-dTAG cells. **j.** Scatterplots like in Supplementary Fig. 5k, but for GANP-2xHA-dTAG cells. **k.** Scatter plot as in Fig. 3c, but of pA^+^ RNA log_2_ fold abundance changes resulting from dTAG^V^-1 treatment of ZFC3H1- (y-axis) or GANP- (x-axis) 2xHA-dTAG cells. **l.** Boxplots of total, nuclear and cytoplasmic pA^+^ RNA log_2_ fold abundance changes after dTAG^V^-1 treatment for 4 hours of GANP-2xHA-dTAG cells of PAXT-sensitive multi-exonic (>4 exons) mRNAs compared to control group of multi-exonic mRNAs. PAXT-sensitive mRNAs with log_2_ fold changes below or above 0.5 in either of the compartments are shown.

**Supplementary Figure 8.**
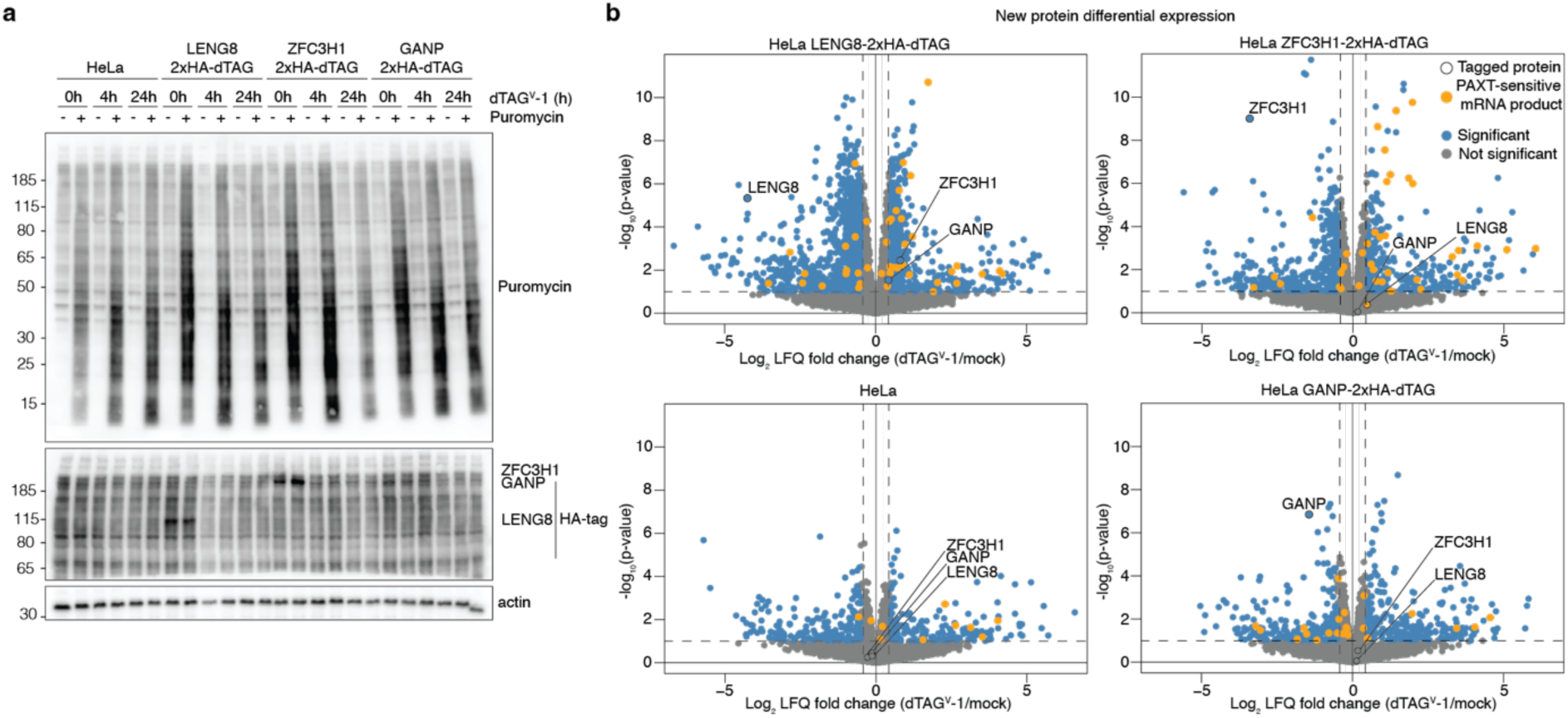
Nuclear pA^+^ RNAs are exported by TREX-2 if not retained and degraded by PAXT. **a.** Western blotting analysis of WCEs from maternal HeLa or ZFC3H1-, LENG8- and GANP-2xHA-dTAG cells mock treated or treated for 4 or 24 hours with dTAG^V^-1. Cells were treated (+), or not (-), with 5ug/ml of puromycin for 30 min before harvest. Membranes were probed with puromycin, HA-tag and actin-specific Abs as indicated. **b.** Scatter plots of differential expression analysis of new proteins synthesized during 24 hours labeling in SILAC medium of LENG8-, ZFC3H1- or GANP-2xHA-dTAG cells and maternal HeLa cells treated or non-treated with 500nM of dTAG^V^-1 for 28 hours. Significantly up- (log_2_ fold change > 0.5) or down-regulated (log_2_ fold change < −0.5) proteins are shown in blue. Protein products of PAXT-sensitive mRNAs are shown in orange. LENG8, ZFC3H1 and GANP are highlighted.

**Supplementary Table 1.**
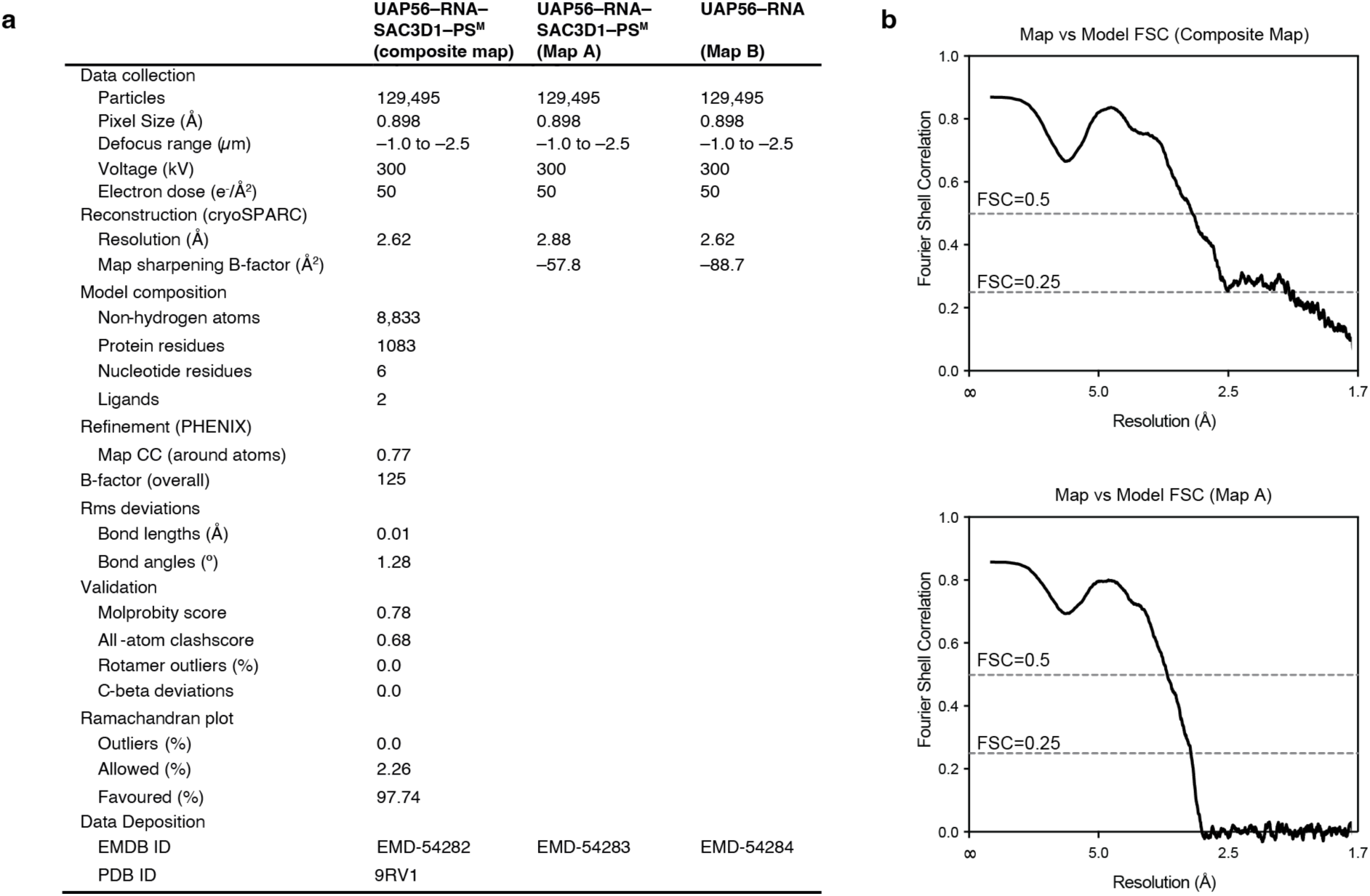
Cryo-EM data collection and refinement statistics.

**Supplementary Table 2 | Results of FLAG immunoprecipitations followed by mass spectrometry.**

**Supplementary Table 3 | Differential expression results.**

**Supplementary Table 4 | Genomic coordinates of sensitive retained and detained introns.**

**Supplementary Table 5 | SILAC whole cell proteomics results.**

**Supplementary Table 6 | Key resources table with vectors, primers, antibodies cell line information.**

